# Stearoyl-CoA Desaturases regulate stem and progenitor cell metabolism and function in response to nutrient abundance

**DOI:** 10.1101/2025.09.17.676943

**Authors:** Kübra B. Akkaya-Colak, Andrea R. Keller, Maria H. Festing, Khushboo Gupta, James Sledziona, Ayobami Adeniyi, Omar West, Patrick Stevens, Maciej Pietrzak, Sue E. Knoblaugh, Helaina Walesewicz, Dominique A. Baldwin, Judith A. Simcox, James Ntambi, Steven K. Clinton, Maria M. Mihaylova

## Abstract

Dietary components and metabolites play a critical role in regulating intestinal stem and progenitor cell function and proliferation. Here we show that Stearoyl-CoA Desaturases (SCDs), which regulate intracellular saturated to monounsaturated fatty acids ratios, are induced in response to nutrient abundance, especially in the distal intestine, and regulate intestinal homeostasis. Genetic or pharmacological inhibition of SCDs altered lipid metabolism, increased ER stress, and reduced proliferative intestinal stem and progenitor cells in intestinal organoids. These effects were largely mitigated by oleic acid supplementation. Intestinal epithelium-specific deletion of *Scd1* and *Scd2* led to metabolic rewiring, leading to expansion of progenitor cell populations. DSS-induced epithelial damage revealed a dependence on SCD enzymes during regeneration, accelerating epithelial damage and inflammation in intestines lacking epithelial *Scd1* and *Scd2*. These findings underscore key metabolic pathways and dependencies that enable intestinal stem and progenitor cells to adapt to nutrient fluctuations and support epithelial tissue regeneration following injury.

## Introduction

The mammalian intestinal epithelium is one of the most rapidly renewing tissues in the body, undergoing complete regeneration approximately every 3-5 days^1^. This dynamic turnover is sustained by intestinal stem cells (ISCs), which reside at the base of crypts and fuel epithelial regeneration through continuous self-renewal and differentiation into secretory and absorptive lineages^2,3^. Given their high proliferative capacity, ISCs and their progenitors have specific metabolic demands that are tightly coupled to nutrient availability and systemic metabolic cues^4,5^.

Recent studies have established that diet and nutrient availability profoundly influence ISC behavior, differentiation, and tissue homeostasis^6–10^. For example, caloric restriction (CR), a reduction in caloric intake without malnutrition, enhances ISC function and self-renewal, in part through inhibition of mTOR signaling^6,11^. Under acute dietary restriction, such as fasting, ISCs adapt by shifting toward fatty acid oxidation, a metabolic reprogramming that boosts their regenerative capacity upon nutrient refeeding^7,12^. In contrast, an excess of lipid or carbohydrate availability, as seen in obesogenic diets, alters ISC function and accelerates adenoma formation^9,13–15^. However, whether high proliferating ISCs and progenitor cells depend solely on exogenous lipids or utilize *de novo* lipogenesis to synthesize fatty acids endogenously remains an open question.

*De novo* lipogenesis is a tightly regulated metabolic process whereby acetyl-CoA is converted to fatty acids, providing essential components for membrane synthesis and signaling lipids^16–18^. Acetyl-CoA carboxylase 1 (ACC1), the rate-limiting enzyme in this pathway, catalyzes the conversion of acetyl-CoA to malonyl-CoA^16^. Recent findings have shown that loss of ACC1 disrupts intestinal crypt formation and compromises organoid viability. Importantly, supplementation with exogenous fatty acids is sufficient to rescue these defects in ACC1-deficient crypts, underscoring the importance of endogenously synthesized lipids in supporting ISC function and maintaining epithelial architecture^19^.

Within the *de novo* lipogenesis pathway, Stearoyl-CoA desaturases (SCDs) introduce a Δ9 double bond into saturated fatty acids (SFAs), generating monounsaturated fatty acids (MUFAs) essential for membrane fluidity and endoplasmic reticulum (ER) homeostasis^20,21^. While SCD1 isoform has been extensively studied in the context of altered cancer metabolism and lipid homeostasis^22,23,24^, the role of SCD1 and SCD2 in the context of intestinal epithelial biology remains largely unexplored. Importantly, how SCD2 is regulated by nutrient availability or systemic signals, and its functional contribution to ISC maintenance and epithelial regeneration, remains unknown.

Here, we investigate the regulation and role of SCD1 and SCD2 in highly proliferating stem and progenitor cells, focusing on how dietary cues modulate their activity within the intestinal epithelium. Our findings provide new insights into the regulation of lipid metabolism in the gut and reveal a critical role for desaturase activity in sustaining ISC function, preserving epithelial homeostasis, and coordinating metabolic adaptation through the balance of fatty acid oxidation and synthesis in nutrient-depleted and nutrient-abundant states. Understanding how metabolism regulates these critical processes may inform future therapeutic strategies across diverse areas, from stem cell biology to metabolism, and offer new avenues for treating tissue damage and gastrointestinal cancers.

## RESULTS

### Nutrient availability controls lipid metabolism in intestinal stem and progenitor cells

Previous studies, including our own, have shown that the small intestine is highly responsive to nutrient availability. Nutrient-sensing pathways, such as mTOR, are tightly regulated during changes in nutrient availability, namely, fasting and feeding^25^ (Supplementary Figure 1A and 1B). To better understand how fasting and refeeding affect intestinal stem cell metabolism, we previously performed RNA-seq on Lgr5-GFP^hi^ intestinal stem cells isolated from short-term fasted (24 hrs.) animals (Supplementary Figure 1C)^7,8^. Strikingly, genes involved in fatty acid oxidation and ketogenesis, such as *Acadl*, *Acot1*, *Pdk4*, and *Hmgcs2*, were significantly upregulated under fasting, while those involved in *de novo* lipogenesis, including *Scd2*, were markedly downregulated (Supplementary Figure 1C). To complement these findings, we performed Western blot analysis of the SCD2 protein in sorted EPCAM+ intestinal epithelial cells from mice fed *ad libitum* or fasted for 24 hours, revealing a significant reduction in SCD2 protein levels under fasting conditions (Supplementary Figure 1D).

Our RNA-seq analysis showed that *Scd2* transcripts were significantly downregulated in intestinal crypt cells in response to fasting. To further validate these findings, we isolated cells from small intestinal and colonic crypts of wild-type male mice subjected to different nutritional states: *ad libitum* feeding or 16 or 24-hour fasting followed by 4 or 24 hours of refeeding (Figure 1A). In contrast to *Scd1* and *Scd2*, expression of key fatty acid oxidation and ketogenesis genes, *Cpt1a* and *Hmgcs2*, increased with fasting and decreased upon refeeding in the small intestinal crypt cells, consistent with previous studies^7,8^ (Figure 1B). Similar pattern was observed in colon crypt cells (Supplementary Figure 1E). Additionally, we found *Scd1*, and especially *Scd2*, to be the predominant isoforms expressed in the intestinal epithelium (Supplementary Figure 1F), consistent with recent reports^26, 27^. Subsequently, we focused our investigations on these two isoforms.

**Figure 1:**
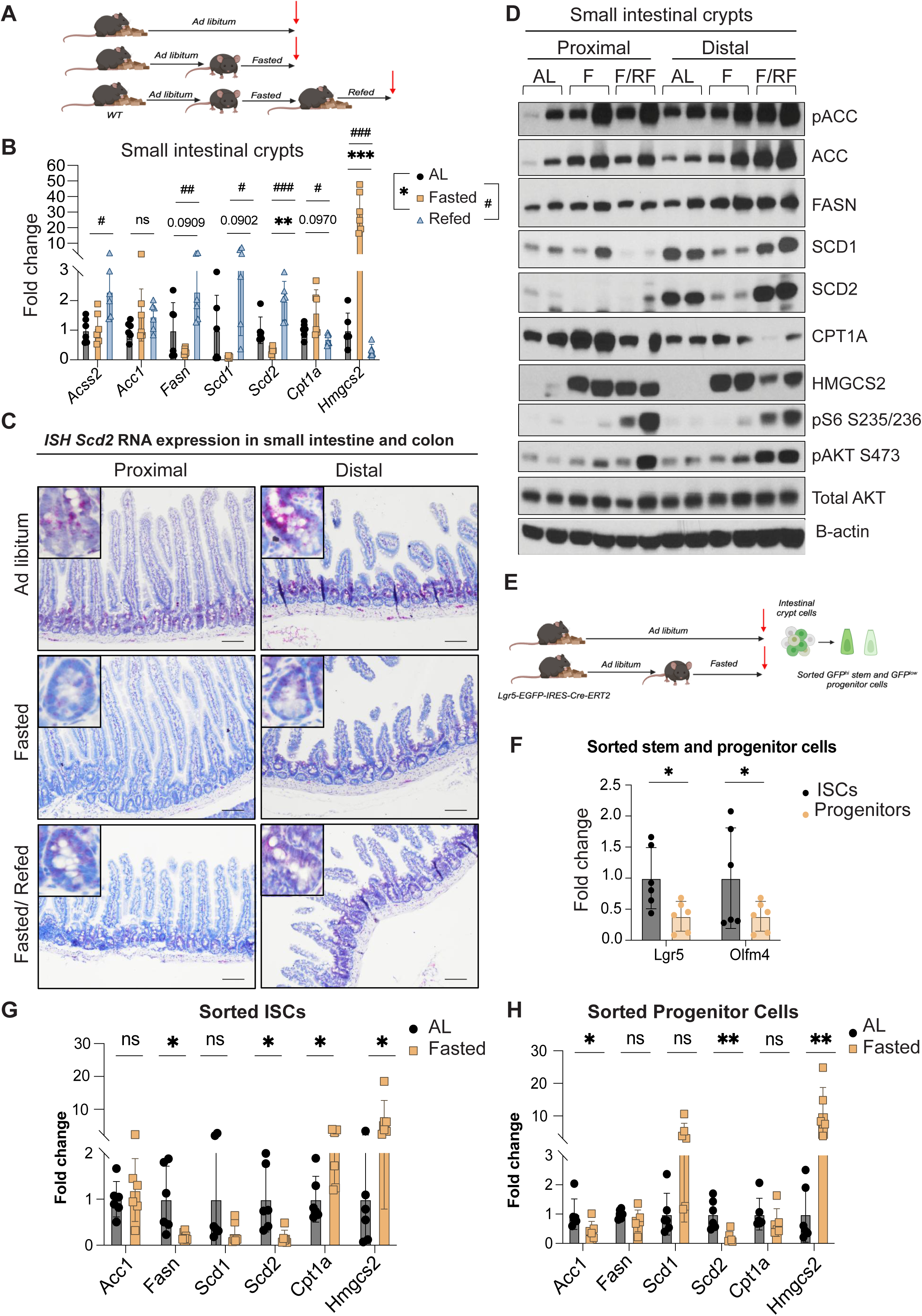
SCD1 and SCD2 levels are regulated by nutrient abundance in intestinal stem and progenitor cells. **(A)** Young WT male animals were set as *ad libitum* (AL), short-term fasted (24 hrs.), and fasted and refed (24 hrs.) animals. Whole small intestinal crypts and colon crypts were isolated from AL, 24 hrs. fasted, and 24 hrs. fasted-refed animals. **(B)** RT-qPCR analysis was performed with small intestinal crypts and colon crypts **(Supp. Figure 1E)** (N=6). **(Supp Figure 1G)** *Scd1* and **(C)** *Scd2* RNA expressions were analyzed by *in situ* hybridization technique in proximal and distal small intestine and colon tissues under different nutrient availability conditions (Images are representative of N=3). **(D)** Western blot analysis was performed to investigate the effect of nutrient availability on SCD1 and SCD2 levels in proximal and distal small intestinal crypts isolated from AL, 16 hrs. fasted, or 16 hrs. fasted followed by 4 hrs. refeeding in WT animals. **(E, F)** *Lgr5+* GFP^hi^ and GFP^low^ small intestinal stem and progenitor cells were isolated from AL and 24 hrs. fasted WT animals by FACS sorting. **(G, H)** RT-qPCR was performed to investigate the expression of *de novo* lipogenesis, fatty acid oxidation, and ketogenesis genes under different nutrient availability (N=6). Data are mean ± SD. *p < 0.05, ***p<0.001, and ****p<0.0005 by Student’s t test, unpaired.

Given that there is metabolic compartmentalization across the intestine, we next asked whether *Scd1* and *Scd2* expression varies across intestinal regions in response to nutrient availability. Proximal and distal small intestine (SI), as well as the colon, were collected to perform *in situ* hybridization analysis. Under nutrient-rich conditions, both *Scd1* and *Scd2* transcripts were highly expressed in the distal end of the small intestine, as well as in the colon, compared to the proximal end of the small intestine. Notably, *Scd2* was the predominant isoform expressed in crypts and transit-amplifying progenitor cells, especially in the distal small intestine and colon. Fasting markedly reduced the expression of both *Scd1* and *Scd2* across all intestinal segments (Figure 1C; Supplementary Figure 1G and 1H). To complement the gene expression analysis, we also evaluated protein levels of SCD1, SCD2, and other key mediators of lipogenesis, fatty acid oxidation, ketogenesis, as well as nutrient-sensing pathways by Western blot (Figure 1D). SCD1 and SCD2 protein levels were reduced in distal crypts during fasting and increased upon refeeding, consistent with our gene expression results. Interestingly there was an enrichment of components of *de novo* lipogenesis pathway in the crypts from distal SI, while key fatty acid oxidation enzyme, CPT1A was enriched in the proximal portion of crypts, suggesting a potential metabolic zonation of the intestine. Others have also previously reported enrichment of FAO program in the proximal end of the intestine^28^.

To investigate the regulation of *Scd* expression specifically in stem and progenitor cells, we isolated GFP^hi^ intestinal stem cells and GFP^low^ progenitors from LGR5-eGFP-CreERT2^+^ mice that were either *ad libitum*-fed or fasted for 24 hours (Figure 1E and 1F). RT-qPCR analysis confirmed a significant reduction in *Scd2* expression in both stem and progenitor cells under fasting (Figures 1G and 1H). Similarly, *Fasn*, an upstream enzyme in the *de novo* lipogenesis pathway, was also downregulated during fasting. Conversely, *Hmgcs2* and *Cpt1a*, key genes involved in ketogenesis and fatty acid oxidation, respectively, were upregulated in fasted stem cells, consistent with our previous findings^7^. Together, these results indicate that SCD1 and SCD2, key enzymes in the *de novo* lipogenesis pathway, are dynamically regulated in response to nutrient availability in high proliferating intestinal stem and progenitor cells.

### Pharmacological inhibition of Stearoyl-CoA desaturase enzymes impairs intestinal stem cell function and induces ER stress

While previous studies have shown that deletion or inhibition of the SCD1 enzyme increases ER stress-induced cell death in various human and murine cancer cells ^21,22^, it remains unclear whether a similar mechanism operates in small intestinal crypt cells. To test this and follow up on our initial observations of nutrient-dependent regulation of SCD enzymes in intestinal stem and progenitor cells, we generated organoids from small intestinal crypts and treated them with pharmacological SCD inhibitors, A939572 and CAY10566. Inhibition of SCD enzymes significantly reduced organoid number, size, and crypt domain formation, which contains all stem and progenitor cells. Overall clonogenicity of SCD-inhibited organoids was significantly lower compared to vehicle-treated controls (Figures 2A and 2B). We additionally treated organoids with the ACC1 inhibitor, ND646, which functions upstream of SCDs in the *de novo* lipogenesis pathway. As recently reported^19^, ACC1 is essential in intestinal homeostasis, and its inhibition completely abrogated organoid formation (Figure 2A). Loss of SCD enzymatic activity also downregulated key intestinal stem cell markers *Lgr5* and *Olfm4* (Figure 2C), indicating reduced stem cell numbers in the organoids. Interestingly, pharmacological SCD inhibition also suppressed *Scd2* and *Acc1* levels (Supplementary Figure 2A).

**Figure 2:**
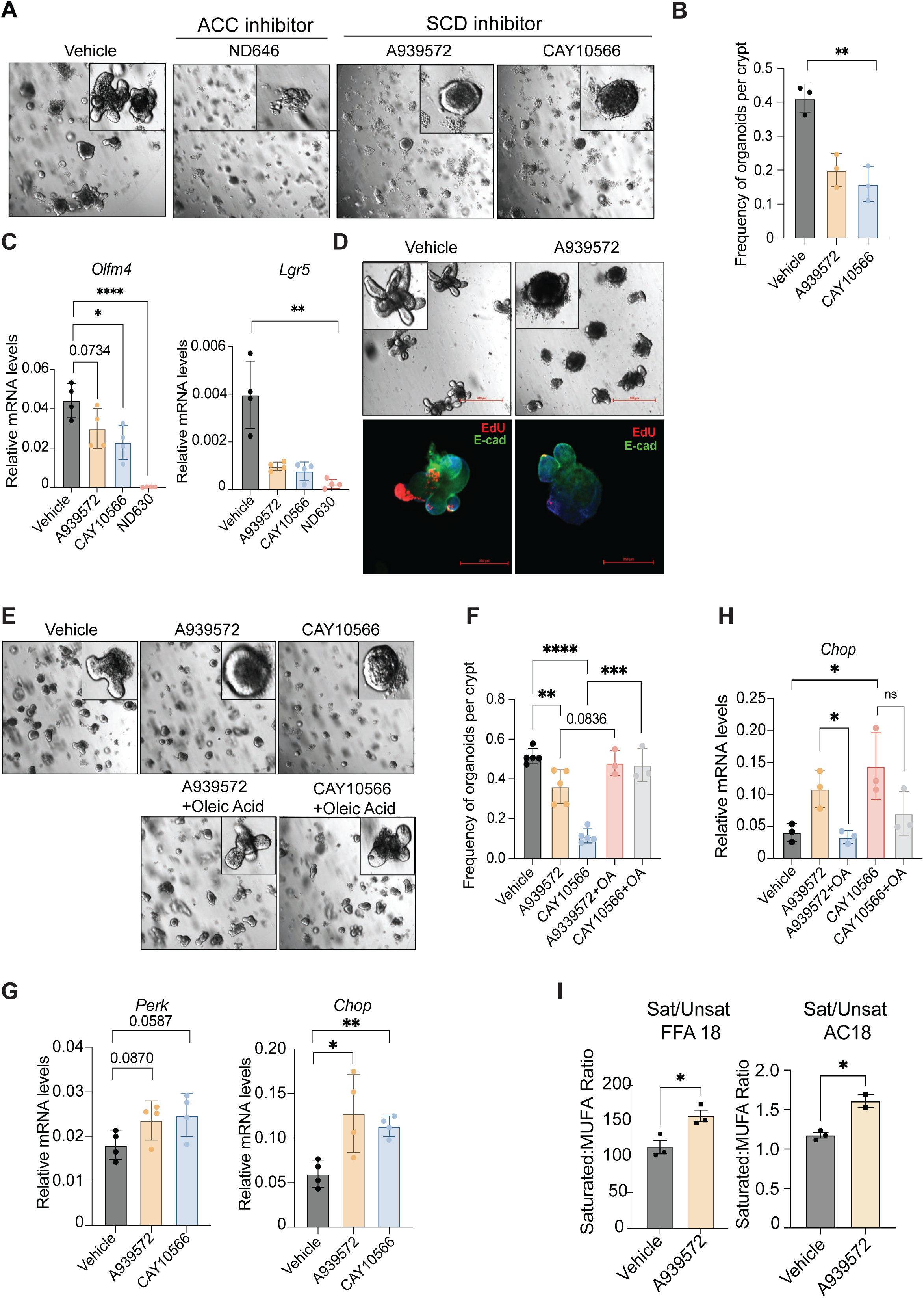
Pharmacological inhibition of SCD1 and SCD2 enzymes reduces stemness and proliferation of intestinal organoids and increases ER stress, which can be mitigated by addition of exogenous MUFAs. The requirement for SCD1 and SCD2 activity in intestinal stem cells was investigated by using mouse organoid models. **(A)** Established WT small intestinal organoids were treated with either vehicle (DMSO), or 250 nM of SCD inhibitors A939572 and CAY10566, or ACC1 inhibitor ND-646 for 96 hrs. (N=4). **(B)** The organoid forming capacity of stem cells and **(C)** the expression levels of intestinal stem cell markers were significantly decreased in inhibitor treated organoids which was investigated by RT-qPCR. **(D)** Organoids were collected and processed for EdU staining which fluorescently labeled proliferative cells. **(E-F)** Exogenous oleic acid addition (50 μM) rescued SCD inhibition (50 nM) phenotype, resulting in increased number of proliferative and differentiated organoids *ex vivo* (N=3-4). While SCD inhibition decreased the number of proliferative cells, addition of oleic acid exogenously rescued the Scd inhibition phenotype. **(G)** Small intestinal organoids were treated with either vehicle (DMSO) or SCD inhibitors A939572 or CAY10566 for four days. RT-qPCR was performed with organoids cDNA to investigate the expression levels of ER stress marker genes. **(H)** RT-qPCR was used to investigate the expression levels of *Chop* in small intestinal organoids under both oleic acid-treated and non-treated conditions, where Scd enzymes were inhibited. **(I)** WT small intestinal organoids were treated with either a DMSO vehicle, or 50 nM of the Scd inhibitor, A939572, for 24 hrs for lipidomics (N=3). Scale bar is 250μm. Graphs and images are representative of 3-4 mice, multiple wells and fields from each biological replicate are counted. Data are mean ± SD. *p<0.05, ***p<0.001, and ****p<0.0005.

To assess the effect of SCD inhibition on intestinal organoid proliferation, we treated organoids with A939572 or vehicle (DMSO), followed by EdU staining. Fluorescence imaging showed a significant reduction in proliferating cells within the crypt domains in SCD-inhibited organoids compared to vehicle-treated ones (Figure 2D), suggesting again impaired stem and progenitor cell proliferation.

To determine if the loss of stem cell function and numbers in SCD1 and SCD2 inhibition is linked to altered MUFA/SFA ratio balance, we performed rescue experiments with oleic acid, one of the main SCD MUFA products. Organoids treated with SCD inhibitors, A939572 and CAY10566, and supplemented with oleic acid, had rescued stem cell proliferation and function, and increased crypt domains like vehicle-treated ones (Figures 2E and 2F). To further explore the cause of reduced proliferation and increased cell death in SCD-inhibited organoids, we examined ER stress markers. RT-qPCR analysis revealed that treatment with A939572 and CAY10566 significantly increased expression of ER stress–induced apoptosis genes, including *Perk* and *Chop* (Figure 2G). Notably, supplementing oleic acid alongside SCD inhibitors reduced *Chop* expression, indicating mitigation of ER stress (Figure 2H).

Finally, we assessed changes in lipid ratios in SCD-inhibited organoids compared to vehicle treated ones. Lipidomic analysis revealed an increase in several SFA, following SCD inhibition, consistent with previous studies in other tissues^29^ (Figure 2I; Supplementary Figure 2B). Together, these findings demonstrate that SCD1 and SCD2 enzymes and their MUFA products are essential to maintain stemness and proliferation in three-dimensional organoid systems. Pharmacological inhibition of SCDs significantly reduces organoid clonogenicity and proliferation, suggesting a dependence on endogenous fatty acid desaturation in this system.

### Loss of *Scd1* and *Scd2* disrupts intestinal stem cell homeostasis and induces ER stress-mediated cell death

To investigate further the role of SCD1 and SCD2 enzymes in intestinal epithelium homeostasis, we generated an intestinal epithelium-specific and inducible *Scd1 (Scd1 iKO*) and *Scd1/2 (Scd1/2 iDKO)* knock out mouse models (Figure 3A) using Villin-CreERT2. Cre recombination was induced by a series of intraperitoneal tamoxifen injections, similar to our prior studies^7^ (Figure 3A). While *Scd1 iKO* and wild-type mice gained weight over time, *Scd1/2 iDKO* mice showed no significant weight gain (Supplementary Figure 3A). Interestingly, deletion of *Scd1* in the intestinal epithelium resulted in a shorter length of the small intestine, while colon length remained unaffected. In contrast, simultaneous deletion of *Scd1* and *Scd2* did not alter small intestinal length significantly but led to a reduction in colon length compared to WT counterparts (Supplementary Figure 3B).

**Figure 3:**
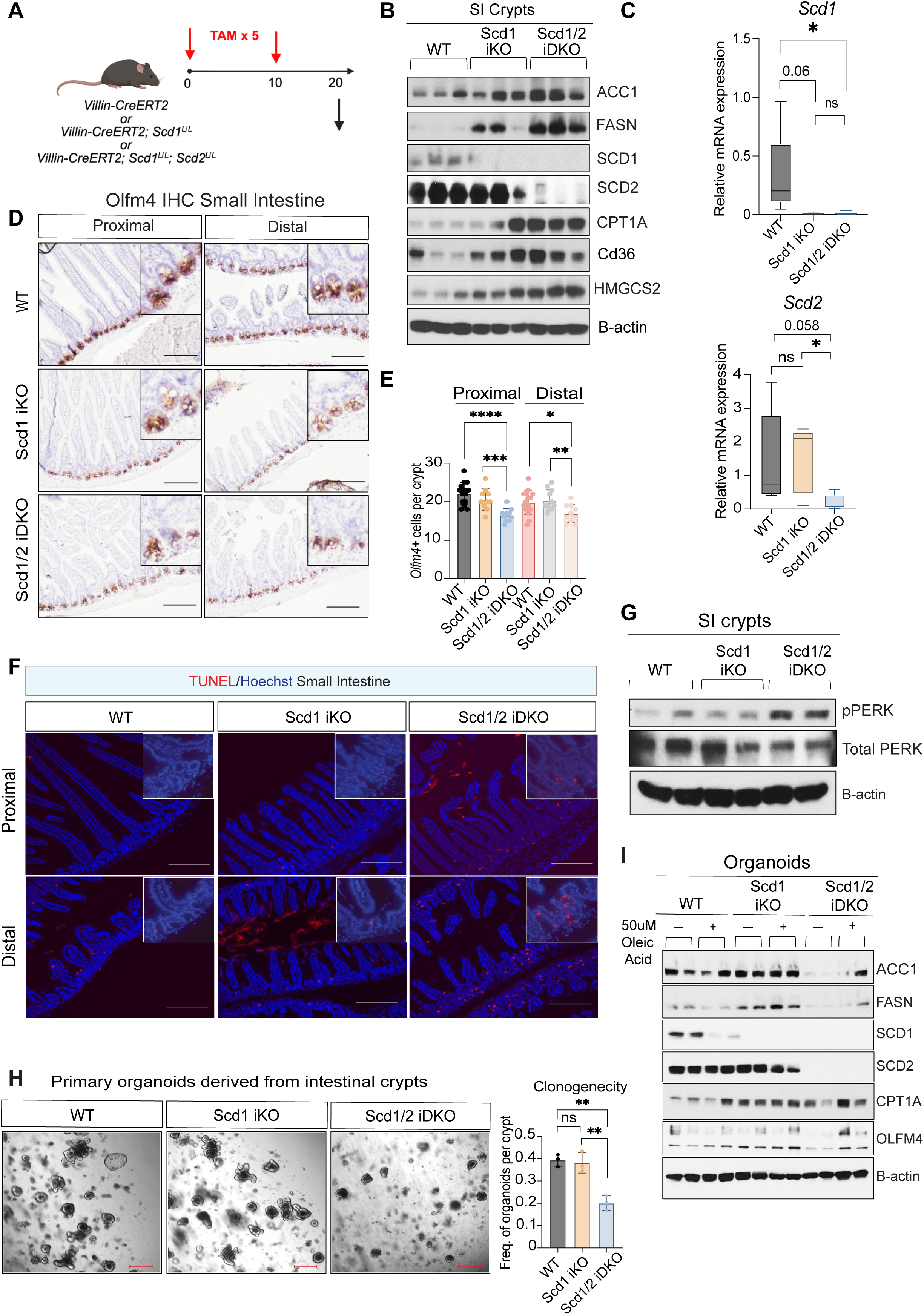
Loss of SCD1 and SCD2 in the intestinal epithelium impairs stem cell function and stemness, inducing ER stress and cell death *in vivo*. **A)** Intestinal epithelium–specific *Scd1* knockout (*Scd1 iKO*) and *Scd1/Scd2* double knockout (*Scd1/2 iDKO*) mice were generated by administering five intraperitoneal tamoxifen injections to male *Villin-CreERT2;Scd1^L/L^; and Villin-CreERT2; Scd1^L/L^; Scd2^L/L^ animals*, respectively. Tamoxifen-injected *Villin-CreERT2* mice served as wild-type (*WT*) controls. **(B)** Small intestinal crypt cells were isolated from each genotype and Western blot was performed to confirm gene deletion and assess regulation of *de novo* lipogenesis, fatty acid oxidation, and ketogenesis pathways (N=3). **(C)** *Scd1* and *Scd2* deletion was further validated by RT-qPCR (N=5–7). **(D)** The effect of *Scd1* and *Scd1/Scd2* deletion on intestinal stem cell stemness was assessed by Olfm4 immunohistochemistry in both proximal and distal small intestinal regions (N=2–3). **(E)** Quantification of Olfm4+ cells per crypt was performed in the proximal and distal small intestine. **(F)** TUNEL staining was conducted to investigate the impact of *Scd1* and *Scd1/2* deletion on cell death within the intestinal epithelium, particularly in the crypts. The deletion of both isoforms led to an increase in dead cells in both the proximal and distal regions of the small intestine, with a more pronounced effect observed in the distal small intestine (N=2-3) **(G)** Biochemical analysis through Western blotting of small intestinal crypts isolated from the experimental animals showed increased phosphorylation of *PERK*, indicating heightened endoplasmic reticulum (ER) stress. **(H)** Small intestinal crypts from *WT*, *Scd1 iKO*, and *Scd1/2 iDKO* animals were isolated and cultured in Matrigel to generate organoids and assess crypt cell function and stemness. Organoid images were captured on Day 4, and clonogenicity was quantified. **(I)** Western blot analysis of Day 4 primary organoids confirmed *Scd1* and *Scd2* depletion and showed altered expression of upstream *de novo* lipogenesis and fatty acid oxidation pathway proteins in *Scd1/2 iDKO* organoids, with or without oleic acid treatment. **(Supplementary** Figures 3F**, 3G)** Primary organoids derived from *WT*, *Scd1 iKO*, and *Scd1/2 iDKO* crypts were cultured for 4 days in complete media with or without 50 µM oleic acid. Clonogenicity and the number of crypt domains (buddings) per organoid were assessed on Day 4. Secondary organoids were generated by passaging primary organoids, and the number of crypt domains per organoid was quantified to evaluate regenerative capacity. Graphs and images represent n=3-7 mice; multiple wells, fields and ≥50 crypts were quantified. Data are presented as mean ± SD. *p < 0.05, ***p < 0.001, ****p < 0.0005.

We next confirmed depletion of SCD1 and SCD2 by Western blot analysis and RT-qPCR from isolated intestinal crypts (Figures 3B and 3C). Interestingly, we noted that loss of SCD1 and SCD2 in intestinal crypts led to increased expression of the fatty acid translocase CD36, suggesting a compensatory mechanism where perhaps cells increase import of exogenous lipids to counteract effects from SCD1/2 loss. In addition, we noted an increase in CPT1A levels, implying a potential increase in fatty acid oxidation. *Scd1/2 iDKO* mice also exhibited symptoms including diarrhea, intestinal distension, crypt and villus elongation (Supplementary Figures 3C-E), suggesting potential lipid malabsorption and activation of epithelial regeneration response.

To assess how loss of *Scd1* or *Scd1/2* impacts intestinal stem and progenitor cells, we performed immunohistochemistry for OLFM4, which is enriched in stem and progenitor cells (Figure 3D). *Scd1/2 iDKO* mice showed a significant reduction in Olfm4+ cells in both proximal and distal small intestine (Figure 3E), while *Scd1 iKO* mice exhibited a modest decrease in Olfm4+ cells in the proximal intestine and no change in the distal end compared to *WT*.

To determine whether the reduced number of Olfm4⁺ stem cells following *Scd* deletion was due to increased cell death, we performed TUNEL staining on small intestinal tissues. An increased number of TUNEL⁺ cells was observed in both proximal and, more prominently, distal crypts of *Scd1/2 iDKO* animals, where *Scd1* and *Scd2* are highly expressed in WT mice (Figure 3F). We next asked whether this cell death is associated with ER stress. Western blot analysis of isolated crypt cells revealed increased phosphorylation of PERK, a key marker of ER stress– mediated apoptosis, specifically in the *Scd1/2 iDKO* mice (Figure 3G).

To compare the regenerative capacity of intestinal stem cells in *Scd1 iKO*, *Scd1/2 iDKO* and *WT* mice, we isolated crypts from these animals and generated organoids. While clonogenicity was preserved in *Scd1 iKO* organoids, *Scd1/2 iDKO* crypts formed significantly fewer organoids per crypt plated while also lacking crypt-like budding domains (Figure 3H). Upon passaging, *Scd1/2 iDKO* organoids had further reduced clonogenicity as they failed to generate secondary organoids or maintain structural integrity observed by decreased number of buds per organoid (Supplementary Figure 3F). Collectively, these findings indicate impaired stem cell function and reduced organoid regenerative capacity.

We next asked whether MUFA depletion contributed to this phenotype; we supplemented the culture medium with oleic acid, the major MUFA product of SCD enzymes. Indeed, oleic acid partially rescued both organoid formation and budding capacity in *Scd1/2 iDKO* cultures (Supplementary Figure 3G). Western blot analysis also showed increased OLFM4 levels in oleic acid-treated *Scd1/2 iDKO* organoids, suggesting a restored number of stem and progenitor cells (Figure 3I). Interestingly, deletion of *Scd1* and *Scd2* significantly reduced levels of ACC1 and FASN, key enzymes in the *de novo* lipogenesis pathway, compared to *WT* organoids. Conversely, CPT1A, a key regulator of fatty acid oxidation, was upregulated in both *Scd1 iKO* and *Scd1/2 iDKO* organoids *ex vivo*, suggesting a metabolic shift toward fatty acid oxidation to meet energy demands or to mitigate intracellular SFA accumulation (Figure 3I).

To further assess more directly the effects of *Scd1 and Scd1/2* deletion on intestinal stem and progenitor cells, we generated mice with *Lgr5+* cell-specific deletion of *Scd1* (*Lgr5-Scd1 iKO*) or both *Scd1* and *Scd2* (*Lgr5-Scd1/2 iDKO*), using Lgr5-EGFP-IRES-Cre-ERT2 allele, and isolated GFP^+^ intestinal stem (*GFP^hi^*) and progenitor (*GFP^low^*) cells by FACS (Figure 4A). Recombination and deletion were confirmed by PCR with specific primers for *Scd1* (Supplementary Figures 4A and 4B) and respective controls (Supplementary Figure 4C), as well as for *Scd2* (Supplementary Figures 4D and 4E) and controls (Supplementary Figure 4F) in sorted Lgr5 positive stem (GFP^hi^) and progenitor (GFP^low^) cells. Western blot analysis of sorted intestinal stem and progenitor cells (Figures 4B and 4C) further confirmed depletion of SCD2 in these cell populations, while SCD1 protein was undetectable due to low protein concentration. To evaluate stem cell function, we performed organoid-forming assays by co-culturing sorted *GFP^hi^* stem cells with *WT* Paneth cells. Strikingly, *Lgr5-Scd1/2 iDKO GFP^hi^* cells showed a severe defect in organoid formation compared to *Lgr5*-*Scd1 iKO* and *WT* cells (*Lgr5-WT*) (Figure 4D). When primary organoids were passaged to assess regenerative capacity, only organoids from *WT* and *Lgr5*-*Scd1 iKO* cells formed branched organoid structures. In contrast, *Lgr5*-*Scd1/2 iDKO* organoids failed to regenerate and exhibited signs of stress and cell death (Figure 4E).

**Figure 4:**
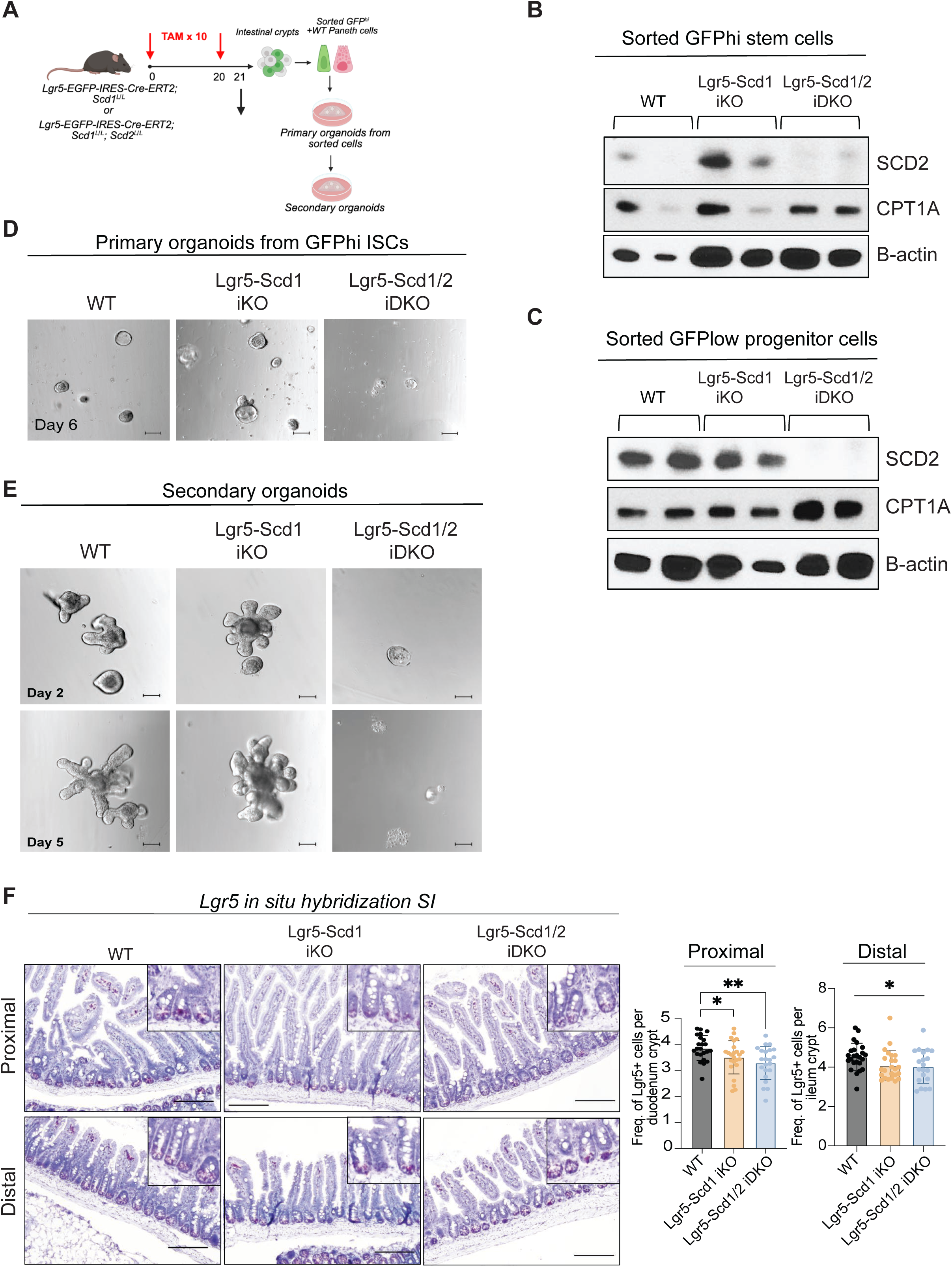
Stem and progenitor cell-specific deletion of *Scd1* and *Scd2* reduces number of *Lgr5+* stem cells and impairs stem cell function. **(A)** Intestinal stem cells (GFP^hi^), progenitor cells (GFP^low^) were FACS sorted from tamoxifen injected *Lgr5-EGF-IRES-Cre-ERT2 (WT)* or *Lgr5-EGF-IRES-Cre-ERT2*, *Scd1^L/L^ (Lgr5-Scd1 iKO)* or *Lgr5-EGF-IRES-Cre-ERT2; Scd1^L/L^; Scd2^L/^ (Lgr5-Scd1/2 iDKO)* animals. **(B, C)** Deletion of *Scd1* and *Scd2* genes was confirmed by Western blot and PCR analysis **(Supplementary Figure 4A-F**) in sorted GFP^hi^ stem and GFP^low^ progenitor cells. **(D)** A mixing experiment was performed with sorted intestinal stem cells from conditional single *Scd1 KO* (*Lgr5*-*Scd1 iKO*) and double *Scd1/2 KO* (*Lgr5*-*Scd1/2 iDKO*) crypts and Paneth cells sorted from *WT* (*Lgr5-EGF-IRES-Cre-ERT2*) animals (N=4). Organoid forming capacity (clonogenicity) of primary organoids was determined on Day 6. **(E)** Secondary organoids were formed by passaging primary organoids. **(F)** Number of cells expressing *Lgr5* gene is shown by *in situ* hybridization in the proximal and distal intestinal epithelium of *WT, Scd1 iKO and Scd1/2 iDKO* animals. Images are representative of 3 or more biological samples and multiple wells, fields and ≥50 crypts were quantified per biological sample. Data are mean ± SD. *p<0.05, ***p<0.001, and ****p<0.0005.

To further examine the impact of stem cell-specific deletion of *Scd1* and combined *Scd1/Scd2* deletion on *Lgr5^+^* intestinal stem cells, we performed *in situ* hybridization for *Lgr5* mRNA in the proximal and distal small intestines (Figure 4F). Our analysis revealed reduction in *Lgr5* positive cells in both *Lgr5*-*Scd1 iKO* and an even more pronounced decrease in the *Lgr5*-*Scd1/2 iDKO* animals, indicating impaired maintenance of Lgr5^+^ stem cells. Together, these findings indicate that SCD1 and SCD2 enzymes are necessary for maintaining intestinal stem cell function and lipid balance. Moreover, imbalance of saturated to monounsaturated fatty acids ratios leads to ER stress, while shift intracellular fatty acid utilization. This may accelerate depletion of intracellular lipid pools, impairing stem and progenitor cell function.

### Deletion of *Scd1* and *Scd2* in the intestinal epithelium induces metabolic rewiring and enhances fatty acid oxidation in small intestinal crypt cells

Next, we investigated the overall impact of *Scd1* and *Scd1/2* deletion in the entire intestinal epithelium. To explore whether *Scd1/2* deletion induces an injury and a metabolic shift, we sacrificed animals at 1-, 3-, or 10-days post tamoxifen induction (Figure 5A). We noted an activation of mTOR signaling, especially at 3- and 10-days post-induction (Figure 5B), as seen by an increase in phosphorylation of T389 S6K, pS6 S235/236, and pS757. Immunohistochemistry further confirmed increased pS6 (S235/236) in both proximal and distal crypts following *Scd1/2* deletion (Figure 5C). mTOR signaling has been shown to promote intestinal stem and progenitor cell growth and facilitate epithelial repair after injury, while disruption of the mTORC1 complex can alter epithelial homeostasis^30^.

**Figure 5.**
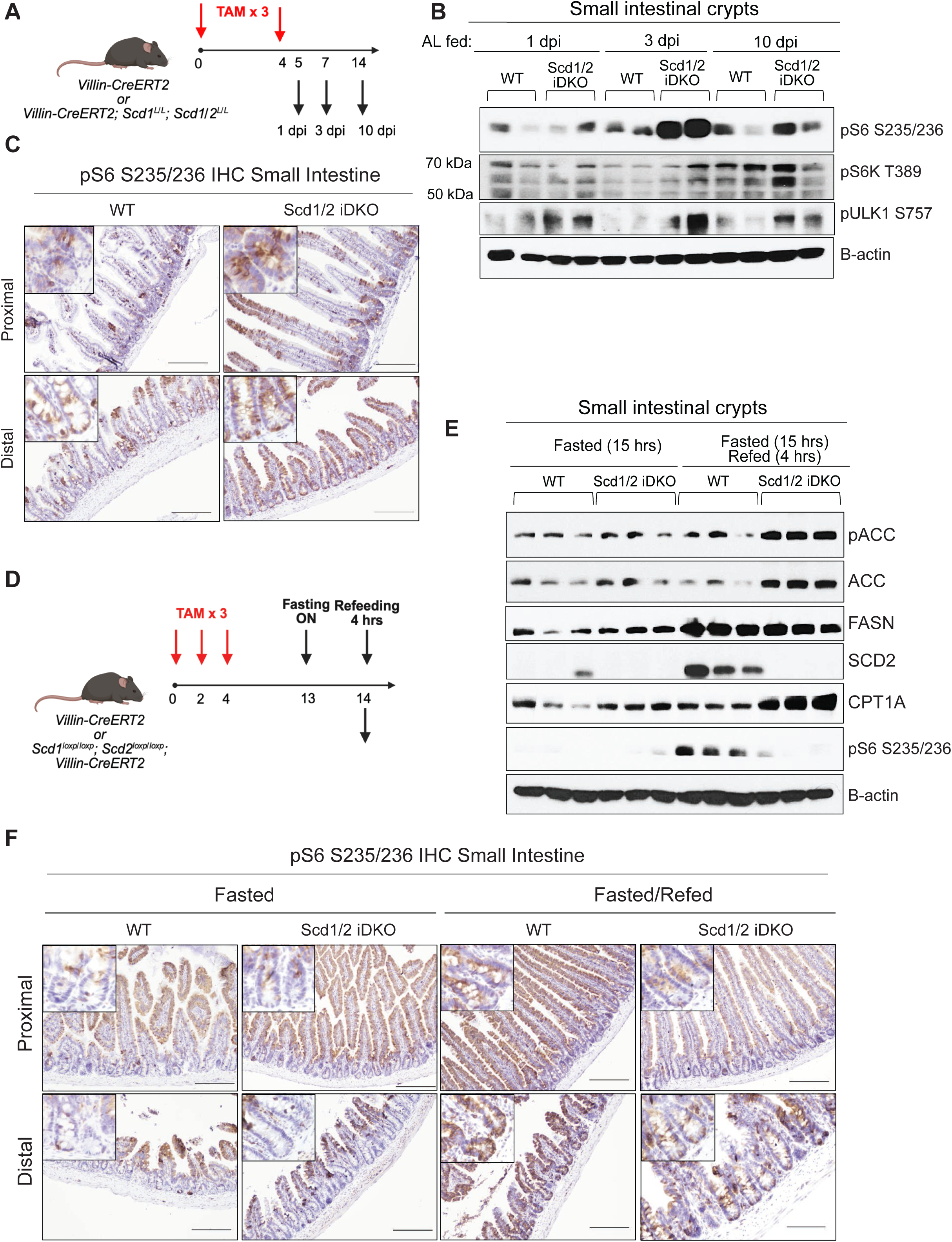
Deletion of *Scd1* and *Scd2* in the intestinal epithelium induces metabolic rewiring via mTOR signaling and promotes fatty acid oxidation in small intestinal crypt cells. **(A)** Schematic of the experimental timeline. Intestinal-epithelium-specific *Scd1/2* double knockout (*iDKO*) mice were generated using *Villin-CreERT2; Scd1^L/L^; Scd2^L/L^* alleles and injected with tamoxifen for three consecutive days. *Villin-CreERT2* animals were also injected as *WT* controls. Mice were sacrificed at 1-, 3-, or 10-days post-injection. **(B)** Western blot analysis of isolated intestinal crypts from *WT* and *iDKO* mice at indicated time points showing increased phosphorylation of S6 (S235/236) and ULK1 (S757), indicative of activated mTORC1 signaling. **(C)** Immunohistochemistry for phospho-S6 (S235/236) in proximal and distal small intestinal sections from *WT* and *iDKO* mice at Day 10 shows increased mTORC1 activity in crypts. **(D)** Schematic of fasting and refeeding experimental timeline. *Villin-CreERT2* and *Villin-CreERT2; Scd1^L/L^; Scd2^L/L^* male animals were i.p. injected 3 times every other day with tamoxifen to generate *WT* and *Scd1/2 iDKO* animals, respectfully. Nine days after the last tamoxifen injection, animals were divided into either fasted or fasted/refed groups (N=3). All animals were fasted overnight for 15 hrs. Only animals in fasted/refed group were refed for 4 hrs. **(E)** Western blot of intestinal crypt lysates showing impaired mTORC1 reactivation in *iDKO* mice after refeeding, as indicated by reduced pS6 (S235/236). Increased levels of Cpt1a and Acc1 are observed in *iDKO* crypts, suggesting a compensatory increase in fatty acid oxidation. **(F)** Representative IHC images of pS6 (S235/236) confirming reduced mTORC1 activation in *iDKO* crypts post-refeeding. Data are representative of at least three biological replicates. Graphs show mean ± SD. *p < 0.05, **p < 0.01, ***p < 0.001.

To further examine nutrient-dependent responses and metabolic flexibility of SCD-depleted mice, we either fasted overnight or fasted and refed for 4-hours prior to analysis (Figure 5D). Interestingly, in contrast to wild-type controls, *Scd1/2 iDKO* crypt cells failed to restore pS6 (S235/236) levels following fasting and refeeding, indicating potentially impaired nutrient transport and mTOR reactivation (Figures 5E and 5F). *Scd1/2* loss also led to elevated CPT1A levels (Figure 5E), particularly following refeeding when it is normally suppressed (Figure 1C). Upstream *de novo* lipogenesis enzymes ACC1 and FASN were also affected, with ACC1 levels increasing in *Scd1/2 iDKO* crypts under fasting-refeeding conditions compared to *WT* (Figure 5E).

We then asked whether the observed metabolic rewiring, along with increased villus length and crypt depth, results from an expansion of fast-proliferating progenitor cell populations. To investigate this, we performed immunofluorescent staining to detect Ki67, a marker of rapidly proliferating cells. Strikingly, in both the proximal and distal small intestine of *Scd1/2 iDKO* animals, we observed a higher number of Ki67⁺ cells compared to their *WT* counterparts (Supplementary Figure 5A).

Next, to trace both Ki67⁺ proliferating cells and their progeny, we injected *WT* and *Scd1/2 iDKO* animals with BrdU for a 24-hour pulse. Immunofluorescent staining for both Ki67 and BrdU was then performed. Interestingly, in the distal small intestine of *Scd1/2 iDKO* animals, there were more BrdU⁺ cells that had migrated along the villi (Supplementary Figure 5B). However, despite this apparent increase in cell migration, the proximal region of *Scd1/2 iDKO* intestines showed a patchy distribution pattern of Brdu^+^ cells.

In summary, these results suggest that SCD1 and SCD2 enzymes are essential for maintaining metabolic and cellular homeostasis in the intestinal crypt cells. Their deletion augments nutrient sensing pathways, leading to metabolic rewiring and altered number and distribution of proliferating cell populations within the intestinal epithelium.

### Deletion of *Scd1* and *Scd2* in the intestinal epithelium exacerbates inflammation and disrupts the intestinal lining in the DSS-induced colitis model

Lipid metabolism plays a key role in regulating intestinal inflammation, with dietary fatty acids capable of either promoting or suppressing inflammatory responses^31^. Oleic acid, a major MUFA produced by SCD enzymes, has been shown to have anti-inflammatory effects in IBD models^32^. Therefore, we hypothesized that deleting *Scd1* and *Scd2* in the intestinal epithelium, key regulators for balancing the MUFA/SFA ratio, would increase inflammation, especially under DSS-induced colitis. To test this, we again induced epithelial-specific deletion of *Scd1* alone or *Scd1*/*Scd2* followed by 2% DSS treatment (Figure 6A). Strikingly, *Scd1/2 iDKO* mice rapidly lost 20% of body weight within 5 days, requiring early study removal (Figure 6B). In contrast, *Scd1 iKO* mice showed no significant weight loss compared to *WT* counterparts. Additionally, colon length was significantly reduced in *Scd1/2 iDKOs* but not in *Scd1 iKOs* (Figures 6C and 6D). *Scd1/2 iDKO* small intestines appeared more severely affected and showed increased crypt cell death relative to *Scd1 iKOs* (Supplementary Figures 6A and 6B), and small intestinal length was reduced in both *Scd1 iKO* and *Scd1/2 iDKO* animals compared to *WT* animals (Supplementary Figure 6C). Colon tissues were submitted for an independent evaluation by a pathologist, which confirmed a significant increase in inflammatory response in the *Scd1 iKO* and *Scd1/2 iDKO* animals compared to *WT* (Figure 6E, 6F). Further histological analysis revealed significant immune infiltration in *Scd1/2 iDKO* colons and increased macrophage infiltration in colon tissues by F4/80 staining (Figure 6G), while *Scd1 iKOs* had milder changes compared to *Scd1/2 iDKOs*. In conclusion, these results indicate that deletion of both *Scd1* and *Scd2* in the intestinal epithelium exacerbates DSS-induced colitis complications and leads to severe inflammation and epithelial destruction in this IBD model.

**Figure 6:**
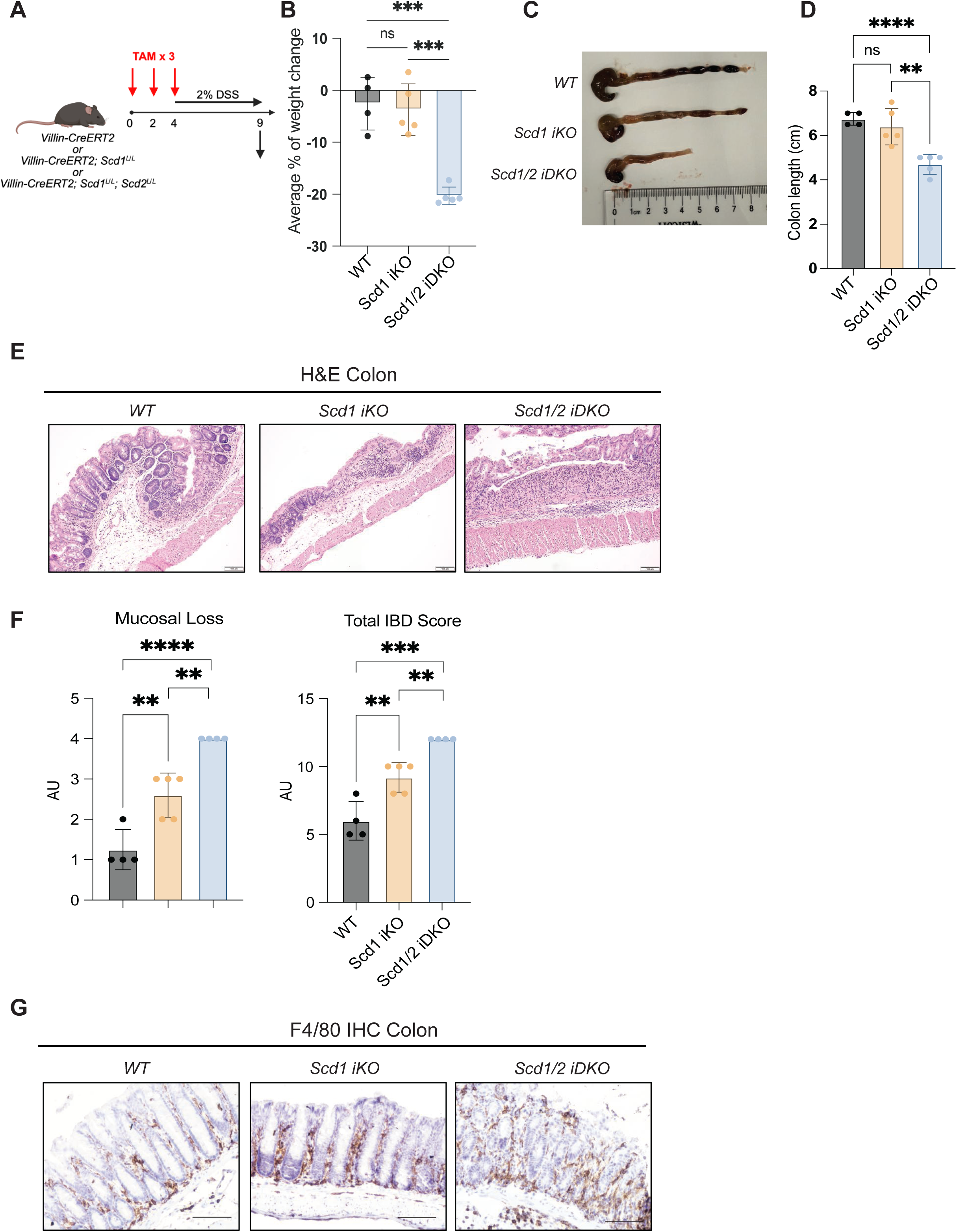
Intestinal epithelium-specific deletion of *Scd1* and *Scd2* exacerbates DSS-induced colitis, leading to severe inflammation, epithelial disruption, and critical weight loss. **(A)** *Villin-CreERT2*; *Scd1^L/L^ (Scd1 iKO)* and *Villin-CreERT2*; *Scd1^L/L^*; *Scd2^L/L^ (Scd1/2 iDKO) male* mice, along with *Villin-CreERT2* controls (*WT*), were administered 3 i.p. tamoxifen injections every other day to induce intestinal epithelium-specific gene deletion. On the day of the final injection, all groups were switched to 2% DSS (dextran sulfate sodium) in drinking water to induce colitis. **(B)** Due to rapid health decline in the *Scd1/2 iDKO* group, all animals (*WT*, *Scd1 iKO*, and *Scd1/2 iDKO*) were euthanized on Day 5. Euthanasia was performed if animals lost ≥20% of their initial body weight (N=4–5 mice per group). **(C-D)** Colon tissues from *WT*, *Scd1 iKO*, and *Scd1/2 iDKO* animals displayed significant shortening in the *Scd1/2 iDKO* group compared to *WT* and *Scd1 iKO.* **(E)** Representative colon images of hematoxylin and eosin (H&E)-stained sections of DSS-exposed animals revealed extensive epithelial damage, crypt loss, and inflammation in *Scd1/2 iDKO* mice. **(F)** H&E-stained colon sections were evaluated by a board-certified veterinary pathologist and scored for inflammation and mucosal loss and given a total IBD Score. **(G)** Immunohistochemistry for F4/80, a macrophage marker, showed elevated immune cell infiltration in the colonic epithelium of *Scd1/2 iDKO* mice, indicating severe inflammation. Data represent mean ± SD. **p < 0.01, ***p < 0.001, ****p < 0.0001.

## Discussion

Diet is a key regulator of adult stem cell function, influencing tissue regeneration, homeostasis, and organismal health^5,33^. We and others have shown that caloric restriction and fasting can enhance stem cell function and delay epithelial aging^6,7^ through suppression of mTORC1 signaling and activation of PPAR-mediated mitochondrial fatty acid oxidation. On the other hand, high-fat or cholesterol-rich diets can promote aberrant intestinal stem cell activity and disrupt epithelial homeostasis^9,34^, yet the dynamics and metabolic rewiring linking dietary inputs to intestinal stem cell (ISC) function are still not fully understood.

While prior studies have assessed how fatty acid oxidation (FAO) impacts ISC function, it is not well known whether high proliferating stem and progenitor cells rely mostly on exogenous fatty acids or engage in *de novo* lipogenesis (DNL). Building on our prior work showing that short-term fasting, followed by refeeding, enhances intestinal stem cell activity via FAO^7^, we now identify Stearoyl-CoA desaturases, SCD1 and SCD2, as critical metabolic regulators of intestinal epithelial homeostasis. We find that these enzymes are dynamically regulated by substrate availability and are essential for maintaining proliferation, self-renewal, and survival of intestinal stem and progenitor cells (Figures 1-4). These findings further uncover the metabolic needs of intestinal stem cells across the gastrointestinal tract and can inform future strategies for epithelial regeneration.

Although extensively studied in cancer, the role of DNL in non-transformed but high-proliferating cells, such as those residing in the intestinal crypt, is not well known. One recent study showed that disruption of ACC1, which facilitates one of the first committed steps in lipid synthesis, impairs crypt architecture and reduces the number of *Lgr5+* ISCs^19^. This phenotype can be rescued by exogenous fatty acids, highlighting the importance of fatty acid metabolism in ISC function. Likewise, obesogenic diets have been shown to rewire ISC differentiation and proliferation, contributing to metabolic diseases and cancer predisposition^15,35^. Our findings in organoid models show that SCD1 and SCD2 play an essential role in sustaining the proliferation and function of intestinal stem and progenitor cells. Mechanistically, these enzymes promote endogenous MUFA synthesis, which supports ISC proliferation and survival by mitigating ER stress-induced cell death (Figure 2). These findings are consistent with prior work in other tissues that show that MUFAs protect against lipotoxic stress^21,22^. Additionally, these enzymes may act on exogenous lipids to desaturate imported SFAs into MUFAs. It has also been recently suggested that ISC and progenitor cells may be more glycolytic, and inhibition of mitochondrial pyruvate metabolism can increase stem and progenitor cell proliferation^36^. Future studies may be needed to examine the utilization of glucose-derived carbons for *de novo* lipid generation in this cell population.

SCD1-mediated DNL plays a key role in tissue and systemic metabolic homeostasis, with dysregulation linked to obesity, inflammation, and cancer^37, 38^. A recent report showed that epithelial-specific deletion of the *Scd1* alters not only intestinal lipid composition but also systemic lipid homeostasis in response to dietary changes^26, 39^. A follow-up study demonstrated further that intestine-specific deletion of *Scd1* modulates systemic energy balance, in part through alterations in bile acid composition and signaling^27^. Utilizing our RNAseq data^7,8^ and complementary approaches, we discovered that *Scd2* transcript and protein are particularly enriched in high proliferating intestinal stem and progenitor cells (Figure 1). Our data further show that intestinal epithelial-specific deletion of *Scd1* and notably *Scd2*, an isoform not previously studied in intestinal stem and progenitor cell homeostasis, alters systemic metabolism, as evidenced by impaired weight gain and reduced adipose tissue in *Scd1/Scd2 iDKO* animals. SCD inhibitors are currently being considered for weight loss management and mitigating obesity driven fatty liver disease^40,41^, and remains to be seen what the long-term consequences will be for prolonged SCD inhibition.

Additionally, we found that intestine-specific deletion of *Scd1* and *Scd2* alters the number and distribution of highly proliferative cells and induces metabolic rewiring *in vivo*. Surprisingly, this is accompanied by activation of mTORC1 signaling, a pathway essential for stem cell proliferation and epithelial regeneration^42^. Recent studies also suggest that fasting and refeeding can impact intestinal regeneration through mTORC1 mediated regulation of protein synthesis^12^.

Following *Scd1/2* loss, cells upregulate the fatty acid translocase CD36, which may lead to higher import of diet-derived lipids. Notably, following overnight fasting and short refeeding, *Scd1/2 iDKO* mice failed to reactivate mTORC1, suggesting delayed responses to nutrient abundance. This was accompanied by compensatory upregulation of FAO genes, indicating a shift toward catabolic metabolism, potentially driven by energy stress and impaired lipogenic homeostasis (Figure 5). Collectively this suggests that SCD enzymes may not be merely downstream responders to nutrient cues but may affect proliferating stem and progenitor cell nutrient sensing.

Previous studies showed that SCD1 is involved in the regulation of inflammation in several tissues, including intestines ^20^. While some studies show global *Scd1* deletion exacerbates DSS-induced inflammation in colitis models^43^, others report minimal impact under controlled conditions^44^. Notably, intestinal *Scd1* loss has been linked to inflammation-driven tumor growth— a phenotype reversed by oleic acid supplementation^45^. Consistent with these findings, our study demonstrates that *Scd1/2* deficiency renders the intestinal epithelium highly susceptible to DSS-induced injury, resulting in severe mucosal loss and heightened immune cell infiltration (Figure 6). Given the known anti-inflammatory properties of MUFAs, particularly oleic acid, our findings suggest that epithelial lipid desaturation is a critical determinant of gut barrier resilience and immune modulation in inflammatory bowel disease. Additionally, mice with colitis may have lower food intake over time and experience calorie and nutrient deficits, potentially revealing a conditional requirement for SCD enzymes during epithelial repair. Future studies will examine whether dietary supplementation with MUFAs can rescue these phenotypes *in vivo*. In parallel, single-cell and spatial transcriptomic analyses, combined with lineage tracing, will be critical to define cell-type–specific responses to impaired lipid homeostasis in stem and progenitor cells, as well as additional lipid-dependent regulatory cues from the niche microenvironment. Collectively, these findings have broad implications for tissue regeneration, aging and disease states such as inflammatory bowel disease, where metabolic plasticity and dietary interventions may be promising therapeutic avenues in the future.

## RESOURCE AVAILABILITY

### STAR METHODS

- **KEY RESOUCES TABLE**
- **EXPERIMENTAL MODEL AND PARTICIPANT DETAILS**

* **MICE**
* **METHOD DETAILS**

## ACKNOWLEDGMENTS

We would like to thank the members of the Mihaylova laboratory for providing critical suggestions and comments on this manuscript. We thank Dr. James Ntambi for kindly gifting the *Scd ^L/L^* and *Scd1^L/L^*; *Scd2^L/L^* animals. This work was supported in part by OSU start-up funds, R00 AG05476, DP2CA271361, the V Foundation Scholar Award and Pew Biomedical Sciences Award. A.K. was supported in part by a Pelotonia Graduate Fellowship. A.K and O.W were supported in part by NIH T32 GM141955. J.S was supported in part by Pelotonia Postdoctoral Fellowship and Cancer Prevention NIH T32 CA2291140-02. H.W. was supported by NSF GRFP (DGE-1747503), D.A.B was supported by the Biology of Aging and Age-Related Diseases T32 (AG0000213). J.A.S is an HHMI Freeman Hrabowski Scholar and had garnered research support from Glenn Foundation and American Federation for Aging Research (A22068 to J.A.S) and an R01 through NIH/NIDDK (R01DK133479 to J.A.S). We thank the Comparative Pathology & Digital Imaging Shared Resource at The Ohio State University Comprehensive Cancer Center, Columbus, OH, for histology support and histopathological evaluation. Images presented in this manuscript were generated using the instruments and services at the Campus Microscopy and Imaging Facility, The Ohio State University (RRID:SCR_025078). These resources are supported in part by grant P30 CA016058, National Cancer Institute, Bethesda, MD. We would like to also thank the shared resource facilities supported by The Ohio State University and the James Comprehensive Cancer Center, as well as the Pelotonia Institute for Immuno-Oncology Immune Monitoring and Discovery Platform staff members, specifically, Kelsi Raynolds, for her expertise and technical support with fluorescence-assisted cell sorting experiments.

## AUTHOR CONTRIBUTIONS

K.B.A.-C. and M.M.M. conceptualized the study. Majority of experiments were conducted by K.B.A.-C with additional experimental support, by A.K, M.F, J.S, A.A, O.W. K.G. A.K helped with RT-qPCR analysis and provided experimental protocols. K.G assisted with Western blots and IHC experiments. M.F. assisted with intestine specific knockout crosses and colony management. J. S. analyzed lipidomics data. J.N. provided the *Scd* floxed animals. H.W., D.A.B, and J.A.S conducted mass spec analysis of SCD-inhibited organoids. S.K provided input and co-mentoring to J.S. S.E.K provided pathology support. M.M.M. supervised experiments. K.B.A.-C and M.M.M wrote the manuscript with additional edits and suggestions by the co-authors.

## Experimental Model and Subject Details

### Mouse Models

All experiments conducted in this study were approved by The Ohio State University Committee on Animal Care (IACUC). All mice in this study were housed at 12 hours light and 12 hours dark cycle with *ad libitum* access to water and food, unless otherwise stated. All mouse strains we used had a C57BL/6 background and obtained from Jackson Laboratory and bred and maintained at the Ohio State University (strain no. 000664).

For fasting and fasting-refeeding experiments, we singly housed young, age-matched wild-type (WT) male animals (12-to 18-week-old) in new cages and removed food completely. Animals were on regular chow diet for their entire life and had access to water all the time. We fasted animals for either 16 or 24 hours. At the end of the indicated time duration mice either were sacrificed (fasting) or given food for additional 4 or 24 hrs (refeeding). *Ad libitum* mice were also singly housed in new cages and had access to food for the full experimental duration.

*Scd^L/L^* animals were a generous gift from Dr. James Ntambi and generated by their lab previously^46^.*Villin-CreERT2* animals were a generous gift from Dr. Sylvie Robine and previously described^47^. *Scd1^L/L^* and *Scd1^L/L^; Scd2^L/L^* mice were crossed to either *Villin-CreERT2* or *Lgr5-EGFP-IRES-CreERT2*^48^ mice to generate *Villin-CreERT2; Scd1^L/L^* or *Lgr5-EGFP-IRES-CreERT2; Scd1^L/L^* and *Villin-CreERT2; Scd1^L/L^; Scd2^L/L^* or *Lgr5-EGFP-IRES-CreERT2; Scd1^L/L^; Scd2^L/L^*. Cre induction was accomplished by administering intraperitoneal (i.p.) tamoxifen injections as indicated. Specifically, animals with a *Villin-CreERT2* background were given 3 or 5 injections, while those with an *Lgr5-EGFP-IRES-CreERT2^+^* background were given 10 injections. Tamoxifen was dissolved in sunflower seed oil to create a final concentration of 10 mg/ml, and 250μl per 25 g of body weight was injected every other day.

### Isolation of small intestine and colon crypts

Intestinal crypt cells were isolated from small intestine as described previously^7^. Colon crypts were also collected for RT-qPCR analysis. Briefly, colon tissues were removed, flushed with ice cold 1X PBS and opened longitudinally. Mucus is then meticulously removed using a microscope slide, and the tissue was washed in ice cold 1X PBS in a 50 mL conical tube, with the PBS being replaced 3 times. The tissue was then cut into approximately 5mm pieces and transferred to a 15 mL conical tube with ice cold 1X PBS plus EDTA (5 mM). The tissue pieces were disrupted by pipetting up and down to release the crypts, settled, and the solution was replaced with fresh ice cold 1X PBS plus EDTA. The tissues were then incubated at 4°C for 15 minutes on a shaker. The PBS/EDTA solution was then carefully removed, and the tissues were washed with cold 1X PBS twice. After ensuring minimal residual PBS, 3 mLs of pre-warmed digestion media, containing colon crypt base media supplemented with 500U of Collagenase IV (Worthington Biochem) per sample, was added to each tube. The tissues are then incubated in a 37°C water bath for 30 minutes to ensure the enzymatic digestion proceeds optimally. To halt the reaction, 10 mLs of cold 1X PBS were added to each tube, and the samples were inverted and pipetted up and down approximately 10 times to release the colon crypts. The solution was transferred to a new 15 mL conical tube, and this process was repeated two more times to ensure a complete crypt isolation. The crypt fractions were then combined, filtered through a 70 μm mesh into a 50 mL conical tube, and were prepared for downstream analyses.

### Organoid culture, drug treatment and rescue experiments

Small intestinal crypts were isolated and embedded in Matrigel (Corning 356231, growth factor reduced) to generate organoids. Small intestinal crypts were cultured in a modified form of crypt culture medium as previously described^7^. Unless otherwise indicated, small intestinal crypts are cultured in crypt media containing Advanced DMEM F12 (Gibco), 2 mM L-Glutamine (Gibco), 100 U/ml Penicillin (Gibco), 1000 ug/ml streptomycin (Gibco), 200 ng/ml Noggin (Peprotech), 50ng/ml R-spondin, 1X B27 (Life Technologies), 3 μM CHIR99021 (Stem Cell Technologies), 1X N2 (Life Technologies), 10 μM Y-27632 dihydrochloride monohydrate (Sigma Aldrich), 1 μM N-acetyl-L-cysteine (Sigma Aldrich), and 15 ng/ml EGF (Peprotech). Small intestinal crypts were encapsulated in 25 μL of Matrigel and plated on a flat-bottom 48-well plate. Then Matrigel droplet with crypts was left to solidify for 15 minutes in a 37°C incubator. 250 μl of complete crypt culture media were added on top of the Matrigel drops with crypts and were incubated at 37°C and 5% CO_2_ in a humidified chamber. Crypt media was refreshed every 3 days.

Acc1 inhibitor ND646 (Cayman) and SCD inhibitors A939572 (Tocris) and CAY10566 (Cayman) were added to crypt culture media at 250 nM concentration and cultured for 3 days at 37°C and 5% CO_2_. On Day 3, crypt media was changed with fresh inhibitors. After 24 hrs, organoids were counted for clonogenicity calculations and images were taken with a light microscope. For rescue experiments, small intestinal crypts were cultured with 50 nM of SCD inhibitors: A939572 (Tocris) and CAY10566 (Cayman) supplemented with 50 μM oleic acid (Sigma). Organoids were cultured 3 days at 37°C and 5% CO_2_. On day 3, crypt culture media was changed with fresh inhibitors and oleic acid. After 24 hrs., organoids were counted for clonogenicity calculations and images were taken with a light microscope.

### RNA isolation and RT-qPCR

The RNA extraction process from intestinal crypts and organoids was performed using Tri Reagent (Life Technologies) and purified with the RNeasy Mini Kit (Qiagen) according to the manufacturer’s instructions. FACS sorted 20,000 *GFP^hi^* (stem cells) and *GFP^low^* (progenitor cells) were dissolved in Tri Reagent solution. Total RNA was isolated sorted stem and progenitor cells using RNeasy Micro Kit (Qiagen). Purified RNA was converted to cDNA using the cDNA Supermix qScript (QuantaBio) and then diluted 1:3 with nuclease free water before amplification with SYBR Green Supermix (Bio-Rad) on Bio-Rad iCycler RT-PCR detection system. The primers used for this process are listed in Table 1.

### *In situ* hybridization

For *in situ* hybridization, small intestine and colon tissues were fixed in 10% formalin, embedded in paraffin, and then sectioned. Single-molecule *in situ* hybridization was performed using the RNAscope 2.5 HD Detection Kit-Red (Advanced Cell Diagnostics) following the manufacturer’s protocol. The sections were first incubated with RNAscope Target Retrieval Reagents for 15 minutes and then with RNAscope Protease Plus Reagent for 30 minutes. Mouse *Mm-Scd1*, mouse *Mm-Scd2* and mouse *Mm-Lgr5* probes were used.

### Histology and immunohistochemistry

Small intestine and colon tissues were fixed in 10% neutral buffered formalin solution as previously described previously in^7, 32^ and sectioned at 5 um thickness. The sectioned tissues underwent antigen retrieval using Borg Decloaker RTU Solution (Biocare Medical) in a pressure cooker (Instant Pot) for 20 minutes and were then incubated with 0.3% H_2_O_2_ for 30 minutes at room temperature. Slides were then blocked with 5% normal donkey serum (Jackson ImmunoResearch) in TBST for 30 minutes, followed by subsequent 15-minute incubations in 5% donkey serum plus avidin/biotin blocking kit (Vector) solutions.

The following antibodies and concentrations were used: rabbit anti-pS6 S235/236 (1:400, CST #2211), rabbit anti-OLFM4 (1:10000, CST #39141) and rabbit anti-F4/80 (D2S9R) (1:200, CST #70076). Donkey anti-rabbit secondary antibody conjugated with biotin (1:400, Jackson ImmunoResearch) was used. This was followed by incubation with the Vectastain Elite ABC Immunoperoxidase detection kit (Vector Labs) and Dako Liquid DAB+ Substrate (Dako) for visualization. Antibody dilutions for immunohistochemistry were made in Common Antibody Diluent (BioGenex). Slides were counterstained with 50% hematoxylin for 1-2 minutes and mounted with Cytoseal XYL (Thermo). For morphological analysis, Hematoxylin and Eosin staining protocol was followed.

### Immunofluorescence

Small intestine tissues were fixed in 10% buffered formalin solution, paraffin embedded and sectioned in 5 um thickness. The sectioned tissues underwent antigen retrieval using Borg Decloaker RTU Solution (Biocare Medical) in a pressure cooker (Instant Pot) for 20 minutes and were then incubated with 0.3% H_2_O_2_ for 30 minutes at room temperature. Then tissue slides were incubated with 0.2% Triton X-100 in 1X PBS (PBST) with 5% normal donkey serum in a humidifying chamber for 1 hr at room temperature. Slides were incubated in primary antibody solution in PBST with 1% donkey serum overnight at 4°C. Primary antibodies rat anti-Brdu [BU1/75 (ICR1)] (1:1000, Abcam ab6326) and rabbit anti-Ki67 (D3B5) (1:400, CST #9129) were used. Alexa Fluor 555 and Alexa Fluor 488 (Invitrogen) secondary antibodies were used, and slides were incubated for 1 hr at room temperature in secondary antibody solution and in a humidifying chamber in the dark. Slides were mounted with Vectashield Vibrance mounting medium after 5 minutes incubation with Hoechst solution with a 1:1000 dilution in 1X PBS. Confocal images were taken with an Olympus FV3000 Multiconfocal microscope or Nikon light microscope.

### TUNEL assay

TUNEL assay was performed with One-step TUNEL In Situ Apoptosis Kit (Red, Elab Fluor® 555, Elabscience, E-CK-A325) following the manufacturer’s specifications. Briefly, slides were deparaffinized and hydrated with varying concentrations of ethanol solutions (100%, 95%, 90%, 80% and 75%) for 3 mins. Then, slides were incubated with 1X Proteinase K solution at 37°C for 20 mins. Next, slides were incubated with fresh labeling working solution including terminal deoxynucleotidyl transferase (TdT) enzyme at 37°C for 60 mins at dark in a humidifying chamber. After washing slides with PBS 3 times 5 mins each, slides were incubated with DAPI solution at room temperature for 5 mins in the dark. Finally, slides were mounted with Vectashield Vibrance mounting medium and stored in dark until imaging with Nikon light microscope.

### Immunoblotting

Crypts and villi were lysed in Cell Lysis Buffer (CST #9803) supplemented with 1 mM PMSF (CST #8553), cOmplete Protease Inhibitor Cocktail (Roche) and PhosSTOP phosphatase inhibitors (Roche). Protein concentrations were determined by BCA assay and normalized to 1 mg/ml concentration. 20-30 µg of normalized lysates were loaded on Novex precast gels (Life Technologies). The following antibodies were used: rabbit anti-Phospho-Acetyl-CoA Carboxylase (Ser79) (1:1000, CST #3661), rabbit anti-Acetyl-CoA Carboxylase (C83B10) (1:1000, CST #3676), rabbit anti-Fatty Acid Synthase (C20G5) (1:1000, CST #3180), rabbit anti-SCD1 (M38) (1:500, CST #2438), mouse anti-SCD2 (H12) (1:500, Santa Cruz #sc-518034), mouse anti-Cpt1a (1:2000, Abcam ab128568), rabbit anti-HMGCS2 (1:2000, Sigma # SAB2107997), rabbit anti-Phospho-S6 Ribosomal Protein (Ser235/236) (1:2000, CST #2211), rabbit anti-Phospho-p70 S6 kinase (Thr389) (D5U1O) (1:500, CST #97596), rabbit anti-Phospho-Akt (Ser473) (D9E) (1:500, CST #4060), rabbit anti-Akt (pan) (C67E7) (1:2000, CST #4691), rabbit anti-Phospho-PERK (Thr980) (16F8) (1:1000, CST #3179), rabbit anti-PERK (C33E10) (1:1000, CST #3192), rabbit anti-OLFM4 (1:10000, CST #39141), rabbit anti-Phospho-ULK1 (Ser757) (D7O6U) (1:500, CST #14202) and mouse anti-beta-actin (1:2000, Santa Cruz sc-47778). Horseradish peroxidase (HRP)-conjugated IgG secondary antibodies (1:4000, CST) were used. Membranes were developed with Pierce™ ECL Western Blotting Substrate (Thermo Fisher).

### EdU Cell Proliferation Assay

Organoids were grown by above mentioned protocol from WT male animals and passaged for every three days. After desired size and budding morphology was maintained, organoids were harvested and incubated with 50 nM SCD inhibitor A939572 (Tocris) for 4 days. After 4 days, organoids were labeled with EdU (10 uM) for 6 hrs (Click-iT EdU Imaging Kit, Thermo Fisher). After EdU labeling, organoids were fixed by 4% PFA/PBS solution for 20 minutes at room temperature (RT). Then, a whole mount organoid staining protocol^49^ was followed to fluorescently stain organoids. Primary mouse anti-E-cadherin (1:400, BD Biosciences BDB610182), secondary Alexa Fluor Donkey anti-mouse 488 antibodies were used. Slides were stained with 1 ug/ml Hoechst 33342 (Thermo Fisher) staining solution for 5 mins at RT and mounted with Vectashield Vibrance (Vector). Organoid images were taken with a Nikon light microscope.

### FACS sorting and mixing experiments

Small intestinal crypts were isolated and dissociated using TrypLE Express for 1 minute and 15 seconds at 37°C. The cells were then suspended in cold SMEM (Life Technologies, 11380-037), centrifuged at 250 g for 5 minutes, and resuspended in cold SMEM with CD31, CD45, TER119, CD24, CD117, and CD326. Cells were incubated in this antibody cocktail for 20 minutes on ice in the dark. After 20 minutes, cells were washed off from the antibody solution with cold SMEM. Afterward, the cells were suspended in SMEM with the viability dye 7-AAD and kept in this solution until they were sorted. We titrated each antibody and developed a cell sorting template specifically for intestinal stem cells (GFP^hi^), progenitor cells (GFP^low^), and Paneth cells with the assistance of the Pelotonia Institute for Immuno-Oncology Center at the Ohio State University. The cells were sorted using the Aurora CS instrument. After collecting each cell type, we mixed 1,500 GFP^hi^ KO or DKO stem cells with 1,500 WT Paneth cells and seeded them with 25 µl of Matrigel. Once the Matrigel had hardened, 250 µl of crypt culture media was added to each well. The plates were then incubated at 37°C with 5% CO2 for 6 days, with media changes every 3 days. Clonogenicity counting and imaging were performed on Day 5-6. On Day 6, organoids were passaged and seeded in fresh Matrigel and media for generation of secondary organoids.

### DSS colitis experiment

For the inflammatory bowel disease model, young male age-matched between 18-21 weeks old *Villin-CreERT2, Scd1^L/L^; Villin-CreERT2 and Villin-CreERT2; Scd1^L/L^; Scd2^L/L^;* animals were injected 3 times every other day with a final tamoxifen solution final concentration of 10 mg/ml, and 250 μl per 25 g of body weight. On the day of last tamoxifen injection, animals were moved to new cages with water bottles containing 2% dextran sulphate sodium (DSS). Animal weights were recorded daily. The end-of-life criteria were determined as a 20% weight loss from the initial weight, and all animals were sacrificed after 5 days of DSS treatment.

### DSS Histopathology and Statistics

The gastrointestinal tract was isolated, and the colon separated from the cecum. Samples of colon, cecum and occasionally mesenteric lymph nodes were placed into 10% neutral buffered formalin. The colon was prepared in a ‘‘Swiss roll’’ technique, routine-processed for paraffin embedding and stained with hematoxylin and eosin.

Hematoxylin and eosin (H&E)-stained colon sections were evaluated by a board-certified veterinary pathologist (SEK) blinded to the experimental groups. Colitis scoring was adapted from previously reported methods^50, 51^ and included scoring the entire colon for the severity of mucosal loss, mucosal epithelial hyperplasia, degree of inflammation and extent of pathology on a 0–4 scale. Scores were summed for a total IBD score. An overall score was submitted for each section evaluated. To compare one variable between two groups, unpaired t-tests were used, or Mann-Whitney tests were used if the data were not normally distributed.

### Organoid small molecule extraction

Analytes collected for targeted analysis were extracted via a two-part metabolomic-lipidomic extraction technique. In brief, organoids were added to ceramic bead tubes, then homogenized in 1000µL of 1:1 trifluoroethanol: water using a Tissuelyser II (Four 40 sec cycles at 30 Hz; cool at 4°C for 5 minutes every 2 cycles). 100µL aliquots of each sample were transferred to separate 1.5 mL centrifuge tubes for further processing. 200µL of 1:1 methanol: ethanol containing 10µL SPLASH II LIPIDOMIX (Avanti #330709) and 249mg d4-succinate were added to each sample. This was followed by the addition of 200µL water, mixed via pipetting, and allowed to incubate at room temperature for 10 minutes. Samples were then pelleted at 16,000xg for 10 minutes and the supernatant for each was transferred to a Captiva EMR-Lipid cartridge (Agilent Technologies #5190-1002). Cartridge extraction was performed using positive nitrogen pressure. The metabolite-containing flowthrough was collected in 1.5mL centrifuge tubes and the cartridge was rinsed twice with 250µL of 2:1:1 water: methanol: ethanol to further collect metabolites. Lipids were eluted into separate 1.5mL centrifuge tubes by washing the cartridge twice with 600µL 2:1 methanol: dichloromethanol. The metabolite and lipid fractions for each sample were dried via SpeedVac. Metabolite samples were reconstituted in 100µL 70:20:10 acetonitrile: water: methanol. Lipid samples were reconstituted in 100µL methanol.

### Organoid metabolite LC/MSMS analysis

Extracts were separated on an Agilent 1290 Infinity II BioLC System using an InfinityLab Poroshell 120 HILIC-Z column (Agilent 683775-924, 2.7µm, 2.1 x 150mm). The column temperature was maintained at 15°C. Mobile phase A was comprised of 20mM ammonium acetate in water (pH 9.3) and 5 µM medronic acid. Mobile phase B was 100% acetonitrile. The mobile phase gradient began with 10% mobile phase A, increased to 22% over 8 min, then increased to 40% by 12 min, 90% by 15 min, then held at 90% until 18 min before re-equilibrating at 10% until min 23. The flow rate was maintained at 0.4 mL/min for the early parts of each run, then increased to 0.5 mL/min from 19.1 min to 22.1 min. Mass spectrometry analysis was performed on an Agilent 6495 C QqQ MS dual AJS ESI mass spectrometer. This method was performed in polarity-switching mode. The gas temperature was kept at 200°C with a flow rate of 14 L/min. The nebulizer was set to 50 PSI, sheath gas temperature at 375°C, and the sheath gas flow at 12 L/min. The Vcap voltage was set at 3000 V, iFunnel high-pressure RF was set to 150V, and the iFunnel low-pressure RF was set to 60V in positive mode. The Vcap voltage was set at 2500V, the iFunnel high-pressure RF was set to 60V, and the iFunnel low-pressure RF was set to 60V in negative mode. A dMRM inclusion list was used to individually optimize fragmentation parameters. Injection volume was set to 1µL.

### Targeted lipid LC/MSMS analysis

Extracts were separated on an Agilent 1290 Infinity II BioLC System using a ZORBAX RRHD Eclipse C18 column (Agilent 959759-902, 1.8 µm, 2.1 x 1500mm). The column temperature was maintained at 40°C. Mobile phase A was comprised of 0.1% acetic acid in water, while mobile phase B was comprised of 90:10 acetonitrile: isopropanol. The mobile phase gradient began with 15% B, increased to 33% at 3.5 min, then increased to 38% at 5.5 min, then increased to 42% B at 7 min, increased to 48% at 9 min, then increased to 65% at 15 min, then increased to 75% at 17 min, then increased to 85% at 18.5, then increased to 95% at 19.5 min, then decreased to 15% B at 21 minutes and held at 15% B until 26 min. Mass spectrometry analysis was performed on an Agilent 6495 D QqQ MS dual AJS ESI mass spectrometer. This method was performed in negative polarity mode. The gas temperature was kept at 290°C with a flow rate of 10 L/min. The nebulizer was set to 35 PSI, sheath gas temperature at 350°C, and the sheath gas flow at 11 L/min. The Vcap voltage was set to 3500 V while the nozzle voltage was set to 1000 V. A dMRM inclusion list was used to analyze specific transitions with a 60-second window total. The injection volume was set to 10µL.

### LC/MSMS data analysis

LC/MSMS raw data were collected in the Agilent native “.d” format. Raw data were analyzed using MassHunter Quantitative analysis (Agilent Technologies). Raw data files have been uploaded to MassIVE Project “Mihaylova_Intestinal_Lipidomics”.

### Quantification and Statistical Analysis

Unless otherwise stated in the figure legends, all experiments reported in this study were repeated at least three independent times. In the main text and figure legends, *N* represents biological replicates. For organoid assays, at least 2–4 wells per group were analyzed, with a minimum of 3–4 biological replicates. Statistical significance was assessed using unpaired Student’s t-test, unless otherwise specified. No samples or animals were excluded from the experiments. Age- and sex-matched young mice (12 to 18 weeks old) were randomly assigned to *in vivo* experiments. All animal experiments were approved by The Ohio State University Institutional Animal Care and Use Committee (IACUC).

**Supplementary Figure 1: *Scd1* and *Scd2* are the predominant isoforms expressed in the intestinal epithelium, particularly in distal crypt cells, and their expression is regulated by nutrient availability (Related to Figure 1**).

**A)** Immunohistochemistry for pS6 (Ser235/236) and **(B)** Western blot analysis show dynamic regulation of the mTOR signaling pathway in the intestinal epithelium following 24 hours of fasting and 24 hours of refeeding post-fasting, with pronounced changes observed in crypt cells. **(C)** RNA-seq analysis of Lgr5-GFP^hi^ intestinal stem cells were isolated by FACS from either ad libitum fed or 24 h fasted animals’ intestinal crypt cells demonstrate nutrient-dependent modulation of lipid metabolism, showing downregulation during fasting and upregulation upon refeeding, including genes in the *de novo* lipogenesis pathway. **(D)** FACS-sorted EPCAM+ intestinal epithelial cells were isolated from *WT* mice following short-term fasting (24 hrs) or fasting (24 hrs) followed by refeeding (24 hrs), to assess nutrient-dependent changes in protein levels in small intestinal crypt cells. **(E)** Whole colon crypts were isolated from AL, 24 hrs fasted, and 24 hrs fasted-refed animals. RT-qPCR analysis was performed with colon crypts (N=6). **(F)** RT-qPCR analysis of *Scd*1–4 isoform expression in crypt and villus epithelial cells isolated from proximal and distal regions of the small intestine in *ad libitum*–fed *WT* male mice (N=4). **(G)** *In situ* hybridization showing *Scd1* mRNA expression in the intestinal epithelium across the proximal and distal small intestine and colon (N=3). **(H)** *In situ* hybridization showing *Scd2* mRNA expression in the colon epithelium (N=3). Data are mean ± SD. *p < 0.05, ***p<0.001, and ****p<0.0005.

**Supplementary Figure 2: Pharmacological inhibition of Scd1 and Scd2 reduces gene expression in small intestinal organoids (Related to Figure 2**).

**(A)** Wild-type (WT) small intestinal organoids were treated for 96 hours with either vehicle (DMSO), SCD inhibitors A939572 (250 nM) and CAY10566 (250 nM), or the ACC1 inhibitor ND-646 (250 nM) (N = 4 biological replicates). Following treatment, organoids were harvested for RT-qPCR analysis to assess gene expression changes. **(B)** Lipidomics analysis was performed with WT small intestinal organoids treated with either a DMSO vehicle, or 50 nM of the Scd inhibitor, A939572, for 24 hrs (N=3 biological replicates). Data are presented as mean ± SD. Multiple wells and views were counted. Data are mean ± SD. p < 0.05, **p < 0.001, ***p < 0.0005.

**Supplementary Figure 3: Deletion of *Scd1* and *Scd2* in the intestinal epithelium leads to morphological alterations and impaired intestinal stem cell function (Related to Figure 3**).

**A)** Initial and final body weights were recorded for *Villin-Cre^ERT^*(*WT*), *Villin-CreERT*; *Scd1^L/L^;* (*Scd1 iKO*) and *Villin-CreERT2; Scd1^L/L^; Scd2^L/L^* (*Scd1/2 iDKO*) mice over a 20-day period. While *WT* and *Scd1 iKO* mice exhibited weight gain, *Scd1/2 iDKO* mice failed to gain weight during this time, indicating a critical role for *Scd1*/*Scd2* in maintaining intestinal homeostasis and overall metabolic health (N=5-6). **(B)** Small intestine and colon lengths were measured in all genotypes, showing significant reductions in *Scd1/2 iDKO* mice (N=9–13). **(C)** Hematoxylin and eosin (H&E) staining was performed on proximal and distal small intestinal sections to assess morphological changes (N=2–3). Quantification of crypt depth **(D)** and villus length **(E)** revealed structural abnormalities in *Scd1/2 iDKO* intestines. **(F)** Primary small intestinal crypts were isolated and passaged to generate secondary organoids; clonogenic capacity was quantified, demonstrating impaired stem cell proliferation and function in *Scd1/2 iDKO* mice. **(G)** A rescue experiment was performed by supplementing cultures with 50 µM oleic acid, which partially restored clonogenicity and number of crypt domains (buddings). Graphs and images are representative of 3-4 mice, multiple wells and fields counted. Data are mean ± SD. *p<0.05, ***p<0.001, and ****p<0.0005.

**Supplementary Figure 4: Deletion of *Scd1* and *Scd2* genes in intestinal stem and progenitor cells was confirmed by PCR (Related to Figure 4**).

*GFP^hi^* and *GFP^low^* intestinal stem and progenitor cells and Paneth cells were isolated by FACS from intestinal crypt cells of tamoxifen-induced young male *Lgr5-EGF-IRES-Cre-ERT2 or Lgr5-EGF-IRES-Cre-ERT2*; *Scd1^L/L^ or Lgr5-EGF-IRES-Cre-ERT2; Scd1^L/L^; Scd2^L/L^* mice (N=4). PCR was performed on 1,000 sorted cells per reaction to detect Cre-mediated recombination of **(A, B)** *Scd1* and **(C)** its controls and **(D, E)** *Scd2* and **(F)** its controls in the respective cell populations.

**Supplementary Figure 5: Intestine-specific deletion of *Scd1* and *Scd2* leads to an expansion of proliferating progenitor cells, potentially promoting regeneration through activated mTOR signaling (Related to Figure 5**).

*Scd1/2 iDKO* animals were generated by crossing *Villin-CreERT2* with *Scd1^L/L^; Scd2^L/L^* alleles and injected with tamoxifen. *Villin-CreERT2* animals were used as *WT* controls. Animals were sacrificed 10 days after the injection. **(A)** Small intestinal tissues were collected, and immunofluorescent staining was performed to detect Ki67 (red) in both the proximal and distal regions (N=3). **(B)** Nine days after the final tamoxifen injection, animals were injected with BrdU (10 mg/mL) and sacrificed 24 hours later. Whole small intestines were collected, and immunofluorescent staining was performed for BrdU (green) and Ki67 (red) (N=3).

**Supplementary Figure 6. Deletion of *Scd* genes increases inflammation and disrupts crypt morphology in the DSS colitis model (Related to Figure 6**).

*Villin-CreERT2* or *Villin-CreERT2; Scd1^L/L^ or Villin-CreERT2; Scd1^L/L^*; *Scd2^L/L^* male mice were i.p. injected 3 times every other day with tamoxifen to generate *WT*, *Scd1 iKO* and *Scd1/2 iDKO* animals, respectfully. On the day of the last tamoxifen injection, mice were given 2% dextran sulfate sodium (DSS) in drinking water for five days to induce colitis. Small intestinal tissues were collected from all group to perform Hematoxylin and eosin (H&E) staining of proximal **(A)** and distal **(B)** parts. (**C**) Total small intestinal lengths were measured (N=4-5). Data are presented as mean ± SD. *p<0.05, ***p<0.001, and ****p<0.0005.

