## Supplementary figures and images for "Stearoyl-CoA Desaturases regulate stem and progenitor cell metabolism and function in response to nutrient abundance"

### Supplemental Figures

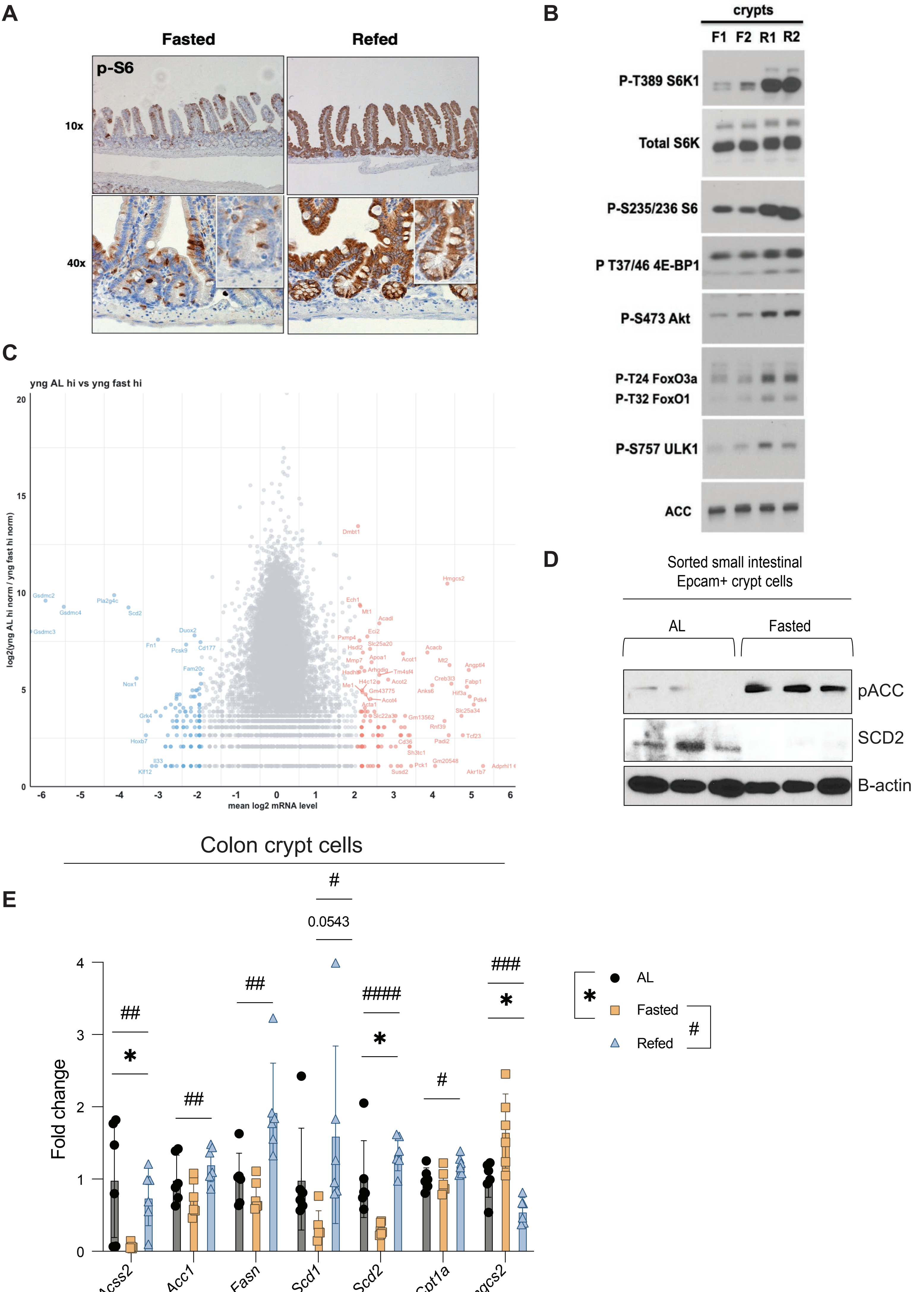

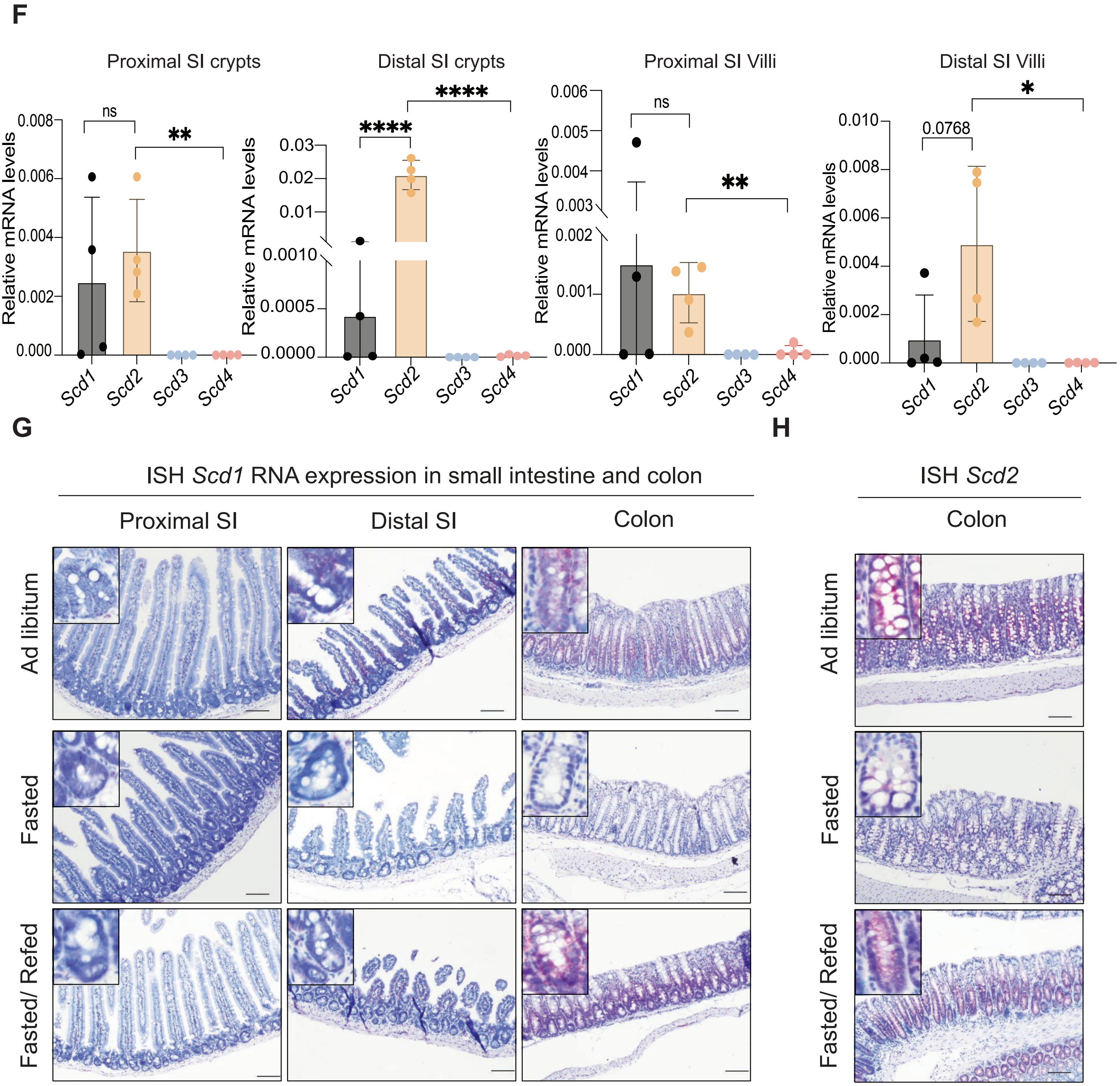

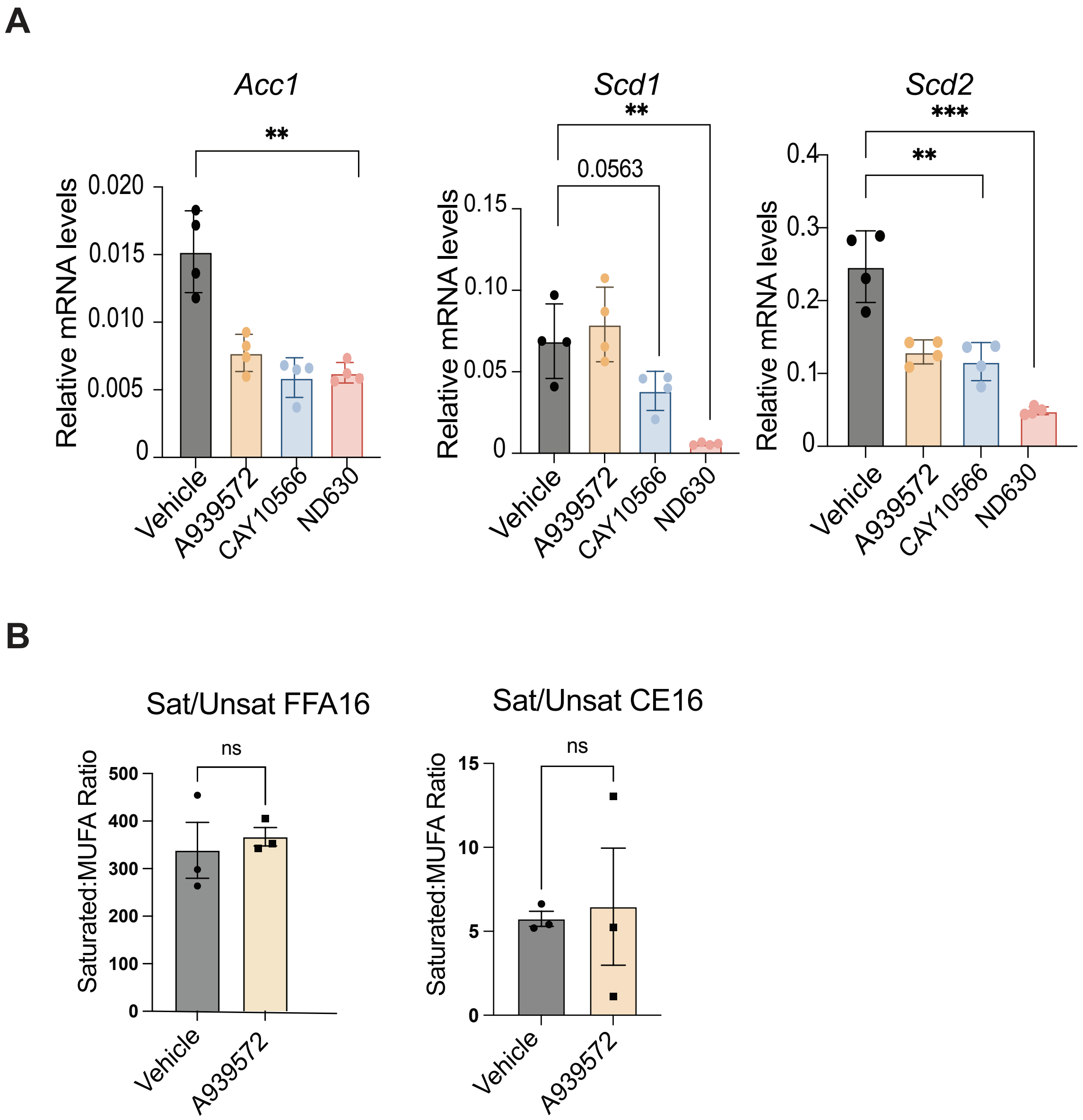

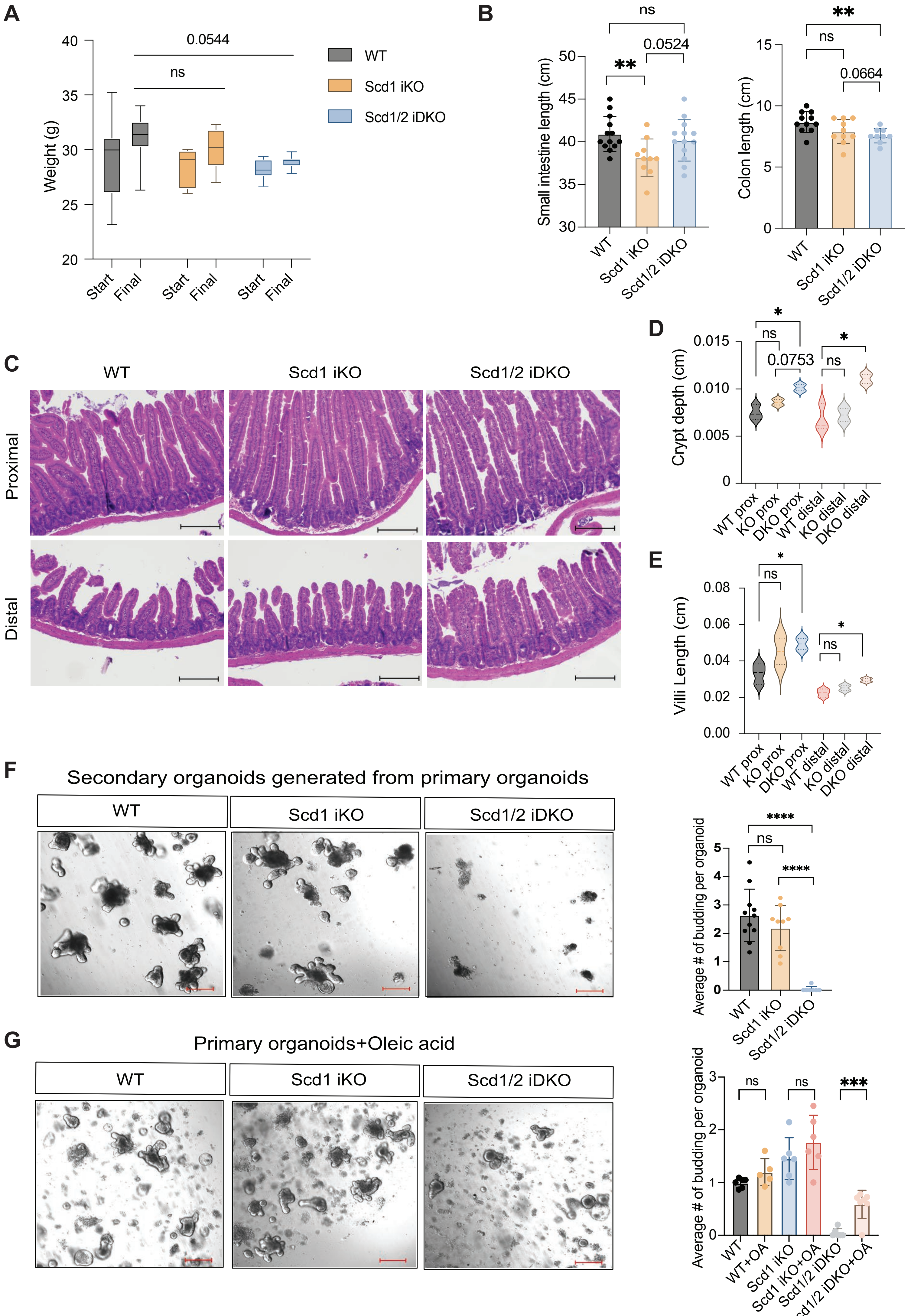

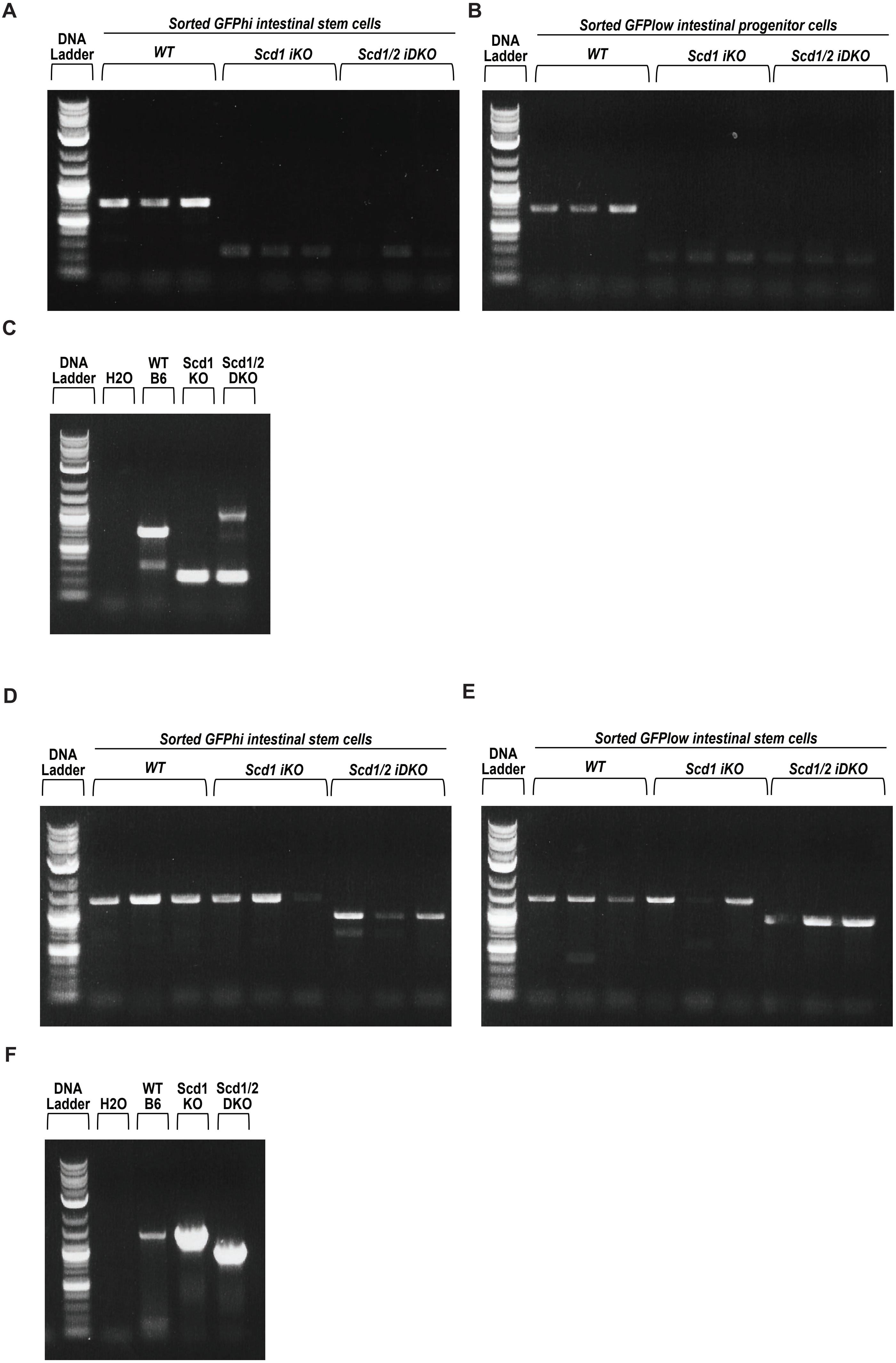

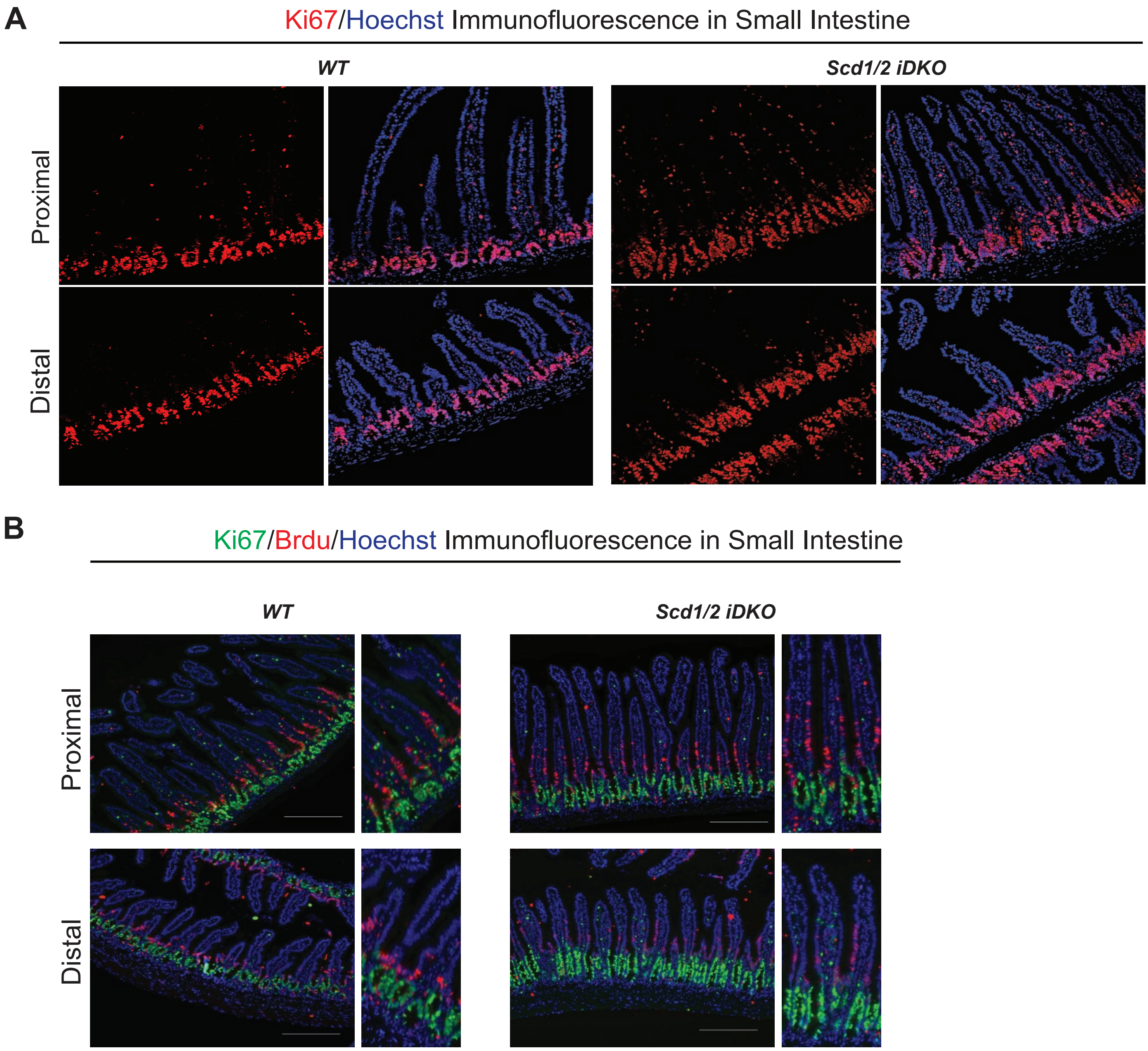

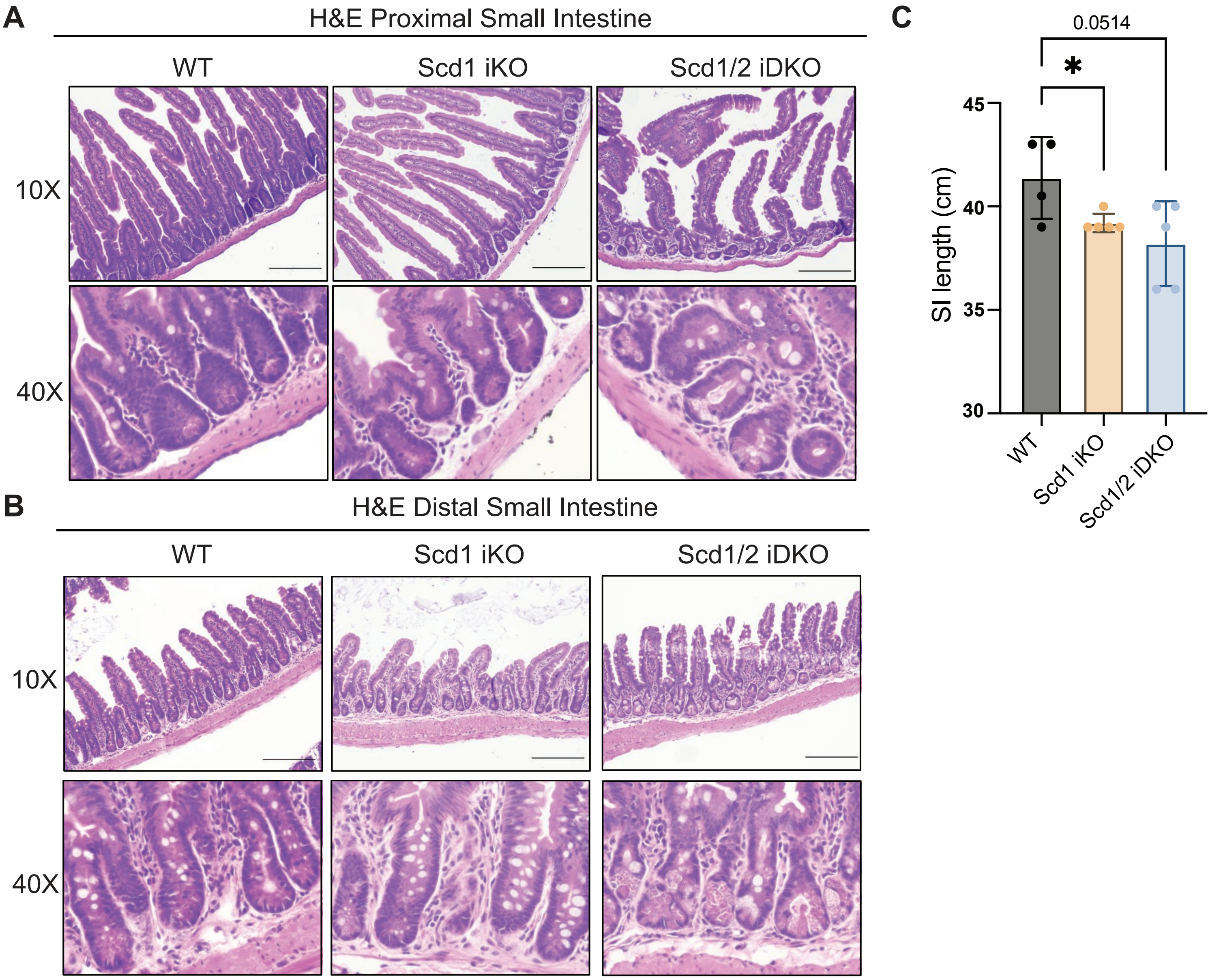
